# Ethanolic leaf extract of *Cleome spinosa* induces apoptosis in Ehrlich ascites carcinoma cells through upregulation of p53 and down-regulation of Bcl-xL in vitro and in vivo

**DOI:** 10.1101/2025.06.11.658929

**Authors:** Priyanka Sharma, Sagarika Ray, Samik Bindu, Tamal Mazumder, Arnab Sen, Subir Chandra Dasgupta, Uday Kishore, Hadida Yasmin

## Abstract

*Cleome* genus is commonly known as spider flower plant or cat’s whiskers, and is the largest genus of the family Cleomaceae, with around 200 species containing a range of remedial applications. Several species of Cleome genus show anti-cancer properties, however, nothing has been reported with genus *spinosa,* an anti-inflammatory and immunomodulatory plant. Thus, the present study examined the likely anti-cancer property of ethanolic leaf extract of *Cleome spinosa* (CSE) in Ehrlich ascites carcinoma (EAC) cells in vitro and in vivo. CSE treatment in vitro significantly caused higher percentage of death in EAC in a dose dependent manner compared to control groups (untreated and ethanol treated). Bright field, fluorescence and electron microscopies revealed apoptotic changes such as surface blebbings, nuclear fragmentation, chromatin condensation and apoptotic body formation, which probably could be the cause of death in EAC cells with CSE treatment. Apoptosis of CSE-treated EAC cells appeared to be mediated through upregulation of pro-apoptotic protein p53 and downregulation of anti-apoptotic protein, Bcl-xL expression, as ascertained through western blot both in vitro and in vivo. CSE was protective toward murine splenocytes by scavenging free radicals; it also reduced the ascitic tumor burden and increased the survival of the tumor-bearing mice compared to the control. Gas Chromatography-Mass Spectrometry (GCMS) analysis identified 43 phytocompounds in CSE. In silico molecular docking analysis indicated that a number of bioactive compounds, such as cycloartenol, stigmasterol, clionasterol, beta-sitosterol, lupenone, and sitostenone that were abundant in the CSE extract, exhibited strong binding affinities for the key apoptosis-regulating proteins, p53 and Bcl-xL. Thus, CSE derived-compounds appear to act as natural antioxidants, and have anti-cancer properties via induction of apoptosis in EAC possibly through upregulation of p53 and downregulation of Bcl-xL proteins in murine model. Hence, CSE could be considered as a promising natural compound for future cancer therapies.

## Introduction

Cancer is the leading cause of mortality, causing 9.7 million deaths; almost 20 million new cases in the year 2020 worldwide ^1–5^ were reported, of which nearly half of the cases occur in Asia, the highest amongst the continents; by 2050 the global burden of cancer is estimated to rise about 77%^6^. Despite considerable progress in the treatment regime of cancer in last few decades, mortalities due to a range of cancer are still high. New target-based FDA (U.S. Food and Drug Administration) approved drugs are continuously being added to the pharmaceutical options ^7,8^. However, due to high manufacturing cost and very less number of truly efficient anti-cancer drugs, the access and effectiveness of experimental drugs remain challenging. Conventional therapies such as radiotherapy and chemotherapy have several side effects and are often immunosuppressive ^9,10^. Thus, there is an argument to explore novel therapeutic approaches which are immunomodulatory, less toxic as well as affordable.

Plants abundantly rich in phenolic compounds, alkaloids, phytosterols, tannins, terpenoids, vitamins, fatty acids and other natural antioxidants have been an important source of medicine for curing different ailment including cancer ^11,12^. According to the WHO, more than 80% of the population in developing countries rely on traditional medicine and indigenous therapies as their age-old practices^13,14^. Pharmaceutical companies are looking for potential anticancer drugs from plant and plant derived compounds due to its cost effectiveness and minimal side effects. Some of these plants derived anti-cancer drugs include curcumin, paclitaxel, adriamycin, vinblastine, and vincristine^15^. Recently much interest has been generated towards plants with immunotherapeutic potential against malignancy such as *Curcuma longa* (Turmeric)^16–19^*, Tinospora cordifolia* (Guduchi) ^20–22^*, Azadirachta indica* (Neem)^23,24^*, Asparagus racemosus* (Shatavari) ^25,26^*, Phyllanthus emblica* (Amla) ^27,28^ *, Albizia lebbeck* (Siris) ^29,30^*, Panax ginseng* (Sansam) ^31,32^*, Moringa oleifera (*Saijna) ^33,34^*, Camellia sinensis (Tea)*^35,36^ and *Withania somnifera* (Ashwagandha) ^37–39^. Several species of Cleome genus have also shown anti-tumor properties. *Cleome* genus is commonly known as spider flower plant or cat’s whiskers and is the largest genus of the family Cleomaceae, with around 200 species containing a range of remedial applications ^40^. Methanol extract of *Cleome arabica* leaf extracts exhibited anti-cancer effects against breast adenocarcinoma, colon carcinoma, neuroblastoma, hepatoma and cervix carcinoma cell lines in vitro^41^. The dichloromethane fraction of the methanolic extract of *Cleome droserifolia* showed 70 to 90% cytotoxicity against MCF-7 (human breast cancer), MDA-MB-231 (triple-negative human breast cancer), and HeLa (human cervical cancer) cell lines ^42^. Five among 18 dammarane-type triterpenes isolated from *Cleome africana* showed potent cytotoxicity against P388 leukemia cells ^43^; methanolic leaf extract of *Cleome gynandra* inhibited Ehrlich ascites carcinoma (EAC) in mice model ^44^. The anti-cancer potential of a few species of genus Cleome, such as *Cleome gynandra, Cleome droserifolia, Cleome africana* and *Cleome arabica* are documented, but nothing has been reported with *Cleome spinosa*.

*Cleome spinosa* is a native of tropical America and is widely cultivated in tropical Asia. It was first reported to get naturalized in West Bengal by Paria in 1980 ^45^. It grows annually during February and March near water lodged areas along road sides, and streams and around forest areas in West Bengal (Fig. 1). These plants are highly tolerant to intense sunlight and drought. Cleome is consumed in most African and South Asian countries due to its high nutritious value. The root and leaf extracts of *Cleome spinosa* exhibited a wide range of antibacterial properties; they are most potent against *Staphylococcus aureus* ^46^. We have shown earlier that ethanolic leaf extract of *Cleome spinosa* (CSE) significantly alleviates the hapten induced delayed type hypersensitivity (DTH) in mouse foot pad by inhibiting TNF-α, suggesting its immunomodulatory property, without demonstrating any histopathological (liver, kidney and spleen) and haematological toxicity ^47^. In another study, intravenous application of CSE, stimulated antibody production in Swiss albino mice when challenged with sheep RBC ^48^. The present study examined the likely anti-cancer properties of CSE, for the first time, in Ehrlich ascites carcinoma (EAC), a spontaneous murine mammary adenocarcinoma. EAC is an undifferentiated carcinoma with wild-type p53 cells with rapid proliferative capacity, and thus, resembles tumors that are highly sensitive to chemotherapy and is also highly metastatic ^49–52^. Here, we carried out in vitro and in vivo studies to understand morphological changes in CSE-treated EAC characteristic of apoptosis, via brightfield, fluorescence and electron microscopy. We also determined the likely mechanism of apoptotic programming in EAC by examining the protein levels of two key apoptosis regulatory proteins, p53 and Bcl-xL. The tumor growth following intravenous CSE treatment in ascites mice was also assessed. GCMS was done to identify the bioactive compounds present in CSE, followed by in silico modeling to identify bioactive molecules exhibiting strong binding affinities to p53 and Bcl-xL.

**Figure 1.**
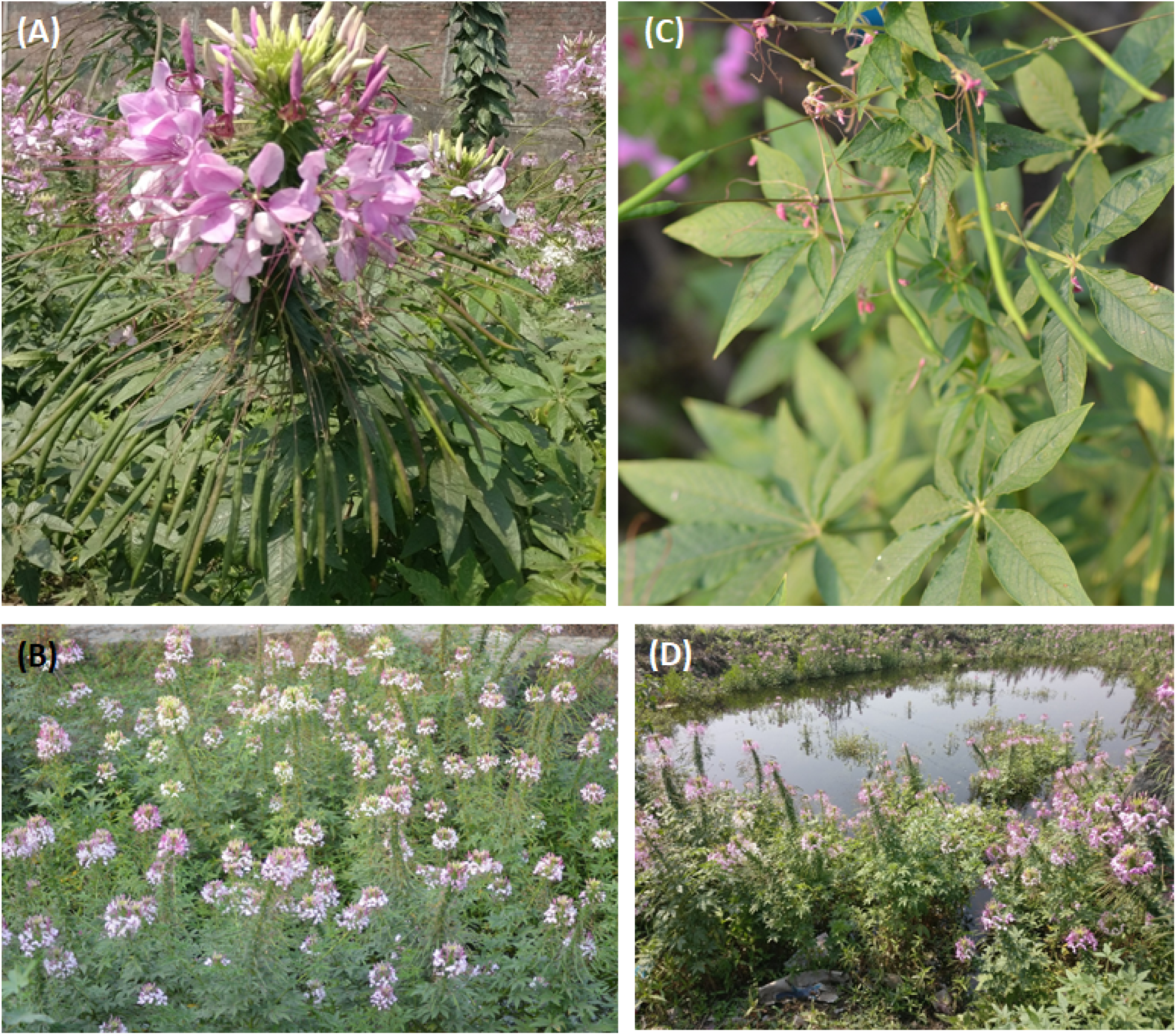
Photographs of *Cleome spinosa* from its collection site. **A)** Cleome inflorescence, often named as spider flower due to appearance of the long, thread-like stamens of the individual flowers and the elongate seedpods that develop below the blooming flowers. **B)** Cleome flowers in full bloom. **C)** Leaves of *Cleome spinosa* used for preparing the ethanolic leaf extract **D)** Presence of *Cleome spinosa* species near wet sites along road side.

## Results

### In vitro CSE treatment induced death of EAC, whereas survival for splenic lymphocytes as observed through trypan blue dye exclusion assay

To ascertain the in vitro doses of CSE that could cause Ehrlich ascites carcinoma cell (EAC) death/apoptosis, the percentage of EAC mortality was compared with the untreated control groups (UNT) at different time points. Two other control groups used were, ethanol (EtOH), the vehicle control as CSE was dissolved in ethanol, and bleomycin (BLM) as a positive control. The efficacy of CSE was compared with UNT, where all the doses of CSE, i.e. 87.6 µg/ml, 146 µg/ml, 182.5 µg/ml and 219 µg/ml caused significantly higher death at 24 h, 48 h and 72 h (Fig. 2Aa). At 72 h, the percentage of death for EAC was 64.4%, 63.9%, 75.4% and 90.4% respectively at 87.6 µg/ml, 146 µg/ml, 182.5 µg/ml and 219 µg/ml dose of CSE, whereas, in control groups, it remained 33.0% (UNT) and 33.4% (EtOH). Among the three doses of BLM (5 µg/ml, 10 µg/ml and 20 µg/ml), 20 µg/ml dose showed the highest rate of EAC cell death being 67.5%, 88.6% and 86.4% at 24 h, 48 h and 72 h respectively (Fig. 2Aa). Figure 2Ab shows the microscopic images of trypan blue stained EAC cells treated with BLM (10 µg/ml and 20 µg/ml) and CSE (146 µg/ml and 182.5 µg/ml) at 24 h, 48 h and 72 h. Compared to control groups (UNT and EtOH) both BLM and CSE treatment exhibited higher number of trypan blue stained EAC cells. We also checked the toxicity of the CSE doses on a normal cell. Thus, in vitro viability of murine splenic lymphocytes was examined through a trypan blue dye exclusion assay with different doses of CSE (Fig. 2B). The lower and moderate doses of CSE, i.e. 87.6 µg/ml, 146 µg/ml and 182.5 µg/ml were not toxic to murine splenocytes (rather stimulatory), showing 61.0%, 56.0% and 38.4% of survival after 48 h, higher than the control (UNT), i.e. 31.5%, but not with the highest dose of CSE (219 µg/ml). The percentage viability of murine splenocytes at 219 µg/ml CSE was lower (11.5%) than the UNT control (31.5%) at the end of 48 h, suggesting 219 µg/ml CSE dose could be lethal for normal cells in vitro (Fig. 2B).

**Figure 2.**
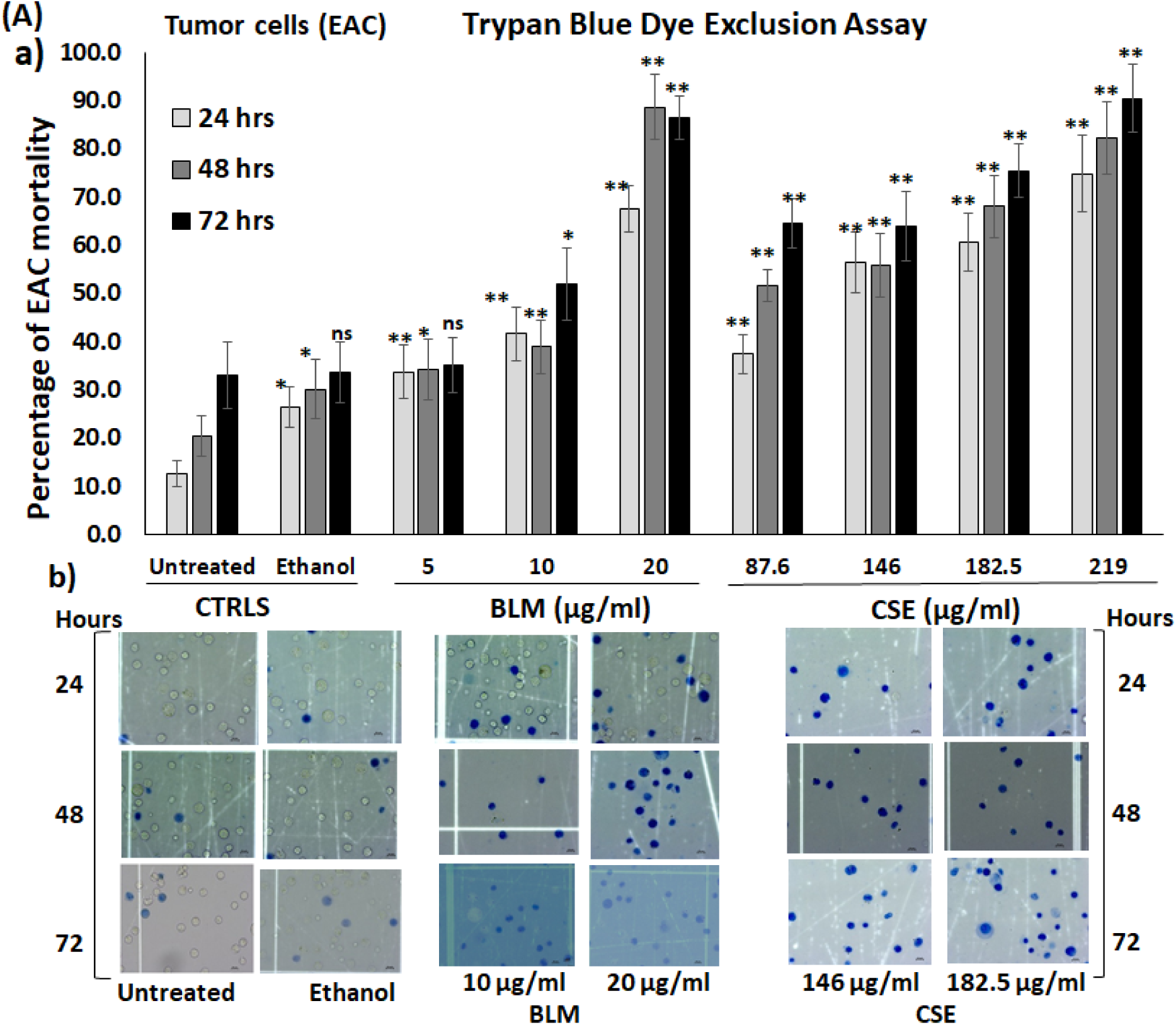
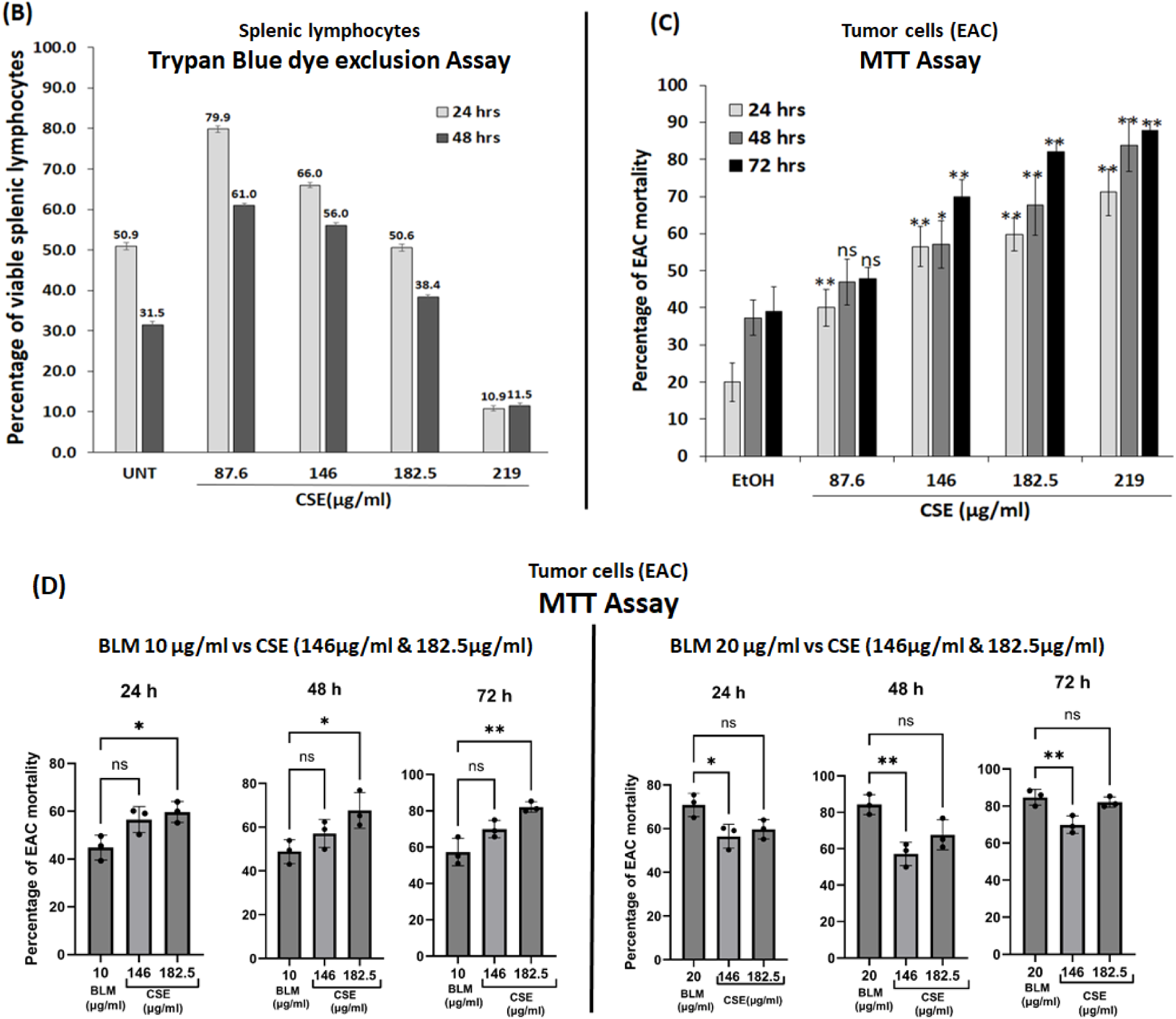
A) In vitro cell mortality of Ehrlich ascites carcinoma cells (EAC) treated with CSE. **a)** Percentage of mortality of EAC through trypan blue dye exclusion test treated with CSE. The statistical significance was calculated comparing the untreated control (UNT) values with other groups; i) ethanol treated (EtOH), as CSE was dissolved in ethanol, ii) BLM (5 µg/ml, 10 µg/ml and 20 µg/ml) used as a positive/standard control, iii) CSE treatment group (87.6 µg/ml, 146 µg/ml, 182.5 µg/ml and 219 µg/ml). Different doses of CSE (87.6 µg/ml, 146 µg/ml, 182.5 µg/ml and 219 µg/ml) induced significantly higher percentage of mortality in EAC cells compared to UNT and EtOH control groups (CTRL). **A) b)** Brightfield microscopy images of trypan blue stained EAC cells (magnification 40X). The microscopic images show high number of trypan blue stained EAC cells with BLM and CSE treatment at 24 h, 48 h and 72 h compared to control groups (UNT and EtOH). **B)** In vitro cell viability of murine splenic lymphocytes treated with CSE was assessed through trypan blue dye exclusion assay. The lymphocytes were treated with different doses of CSE (87.6 µg/ml, 146 µg/ml, 182.5 µg/ml and 219 µg/ml) for 24 h and 48 h. Dose dependent survivality of splenic lymphocytes were observed, where lower doses of CSE were conducive as well as stimulatory and the highest dose i.e. 219 µg/ml was found to be toxic. **C)** Percentage of mortality of EAC with CSE treatment was compared to EtOH through MTT assay. 146 µg/ml, 182.5 µg/ml and 219 µg/ml doses showed significantly high mortality compared to EtOH. **D)** Percentage of mortality of EAC with CSE treatment was compared to BLM (10 µg/ml and 20 µg/ml) through MTT assay. Efficacy of inducing mortality in EAC, 10 µg/ml BLM dose was comparable to 146 µg/ml of CSE and 20 µg/ml BLM was comparable to 219 µg/ml of CSE. Data represent the means ± standard deviation of triplicate values. Significant results were marked with an asterisk (\**P* < 0.05, \*\**P* < 0.01, \*\*\**P* < 0.001 and ns; insignificant).

### In vitro death of EAC with CSE treatment were comparable to Bleomycin (BLM) through MTT assay

The percentage of mortality of EAC with CSE treatment using MTT assay followed the similar pattern of death as observed in trypan blue dye exclusion assay. Various doses of CSE showed significantly high death compared to EtOH at different time points of in vitro treatment. After 72 h, the percentage mortality with CSE was 69.9% with 146 µg/ml, 82.1% with 182.5 µg/ml and 87.7% with 219 µg/ml, whereas EtOH was 39.1% (Fig. 2C). The IC_50_ values of CSE for EAC cells were 128.4 µg/ml, 108.2 µg/ml, and 88.8 µg/ml respectively after 24 h, 48 h and 72 h. The percentage of death of EAC with CSE treatment was also compared with BLM (10 µg/ml and 20 µg/ml). Two CSE doses (146 µg/ml and 182.5 µg/ml) were chosen on the basis of its efficacy to induce EAC death (Fig. 2A and 2C). In spite of showing highest death of EAC, i.e. 87.7% at 72 h, 219 µg/ml dose of CSE was not considered due to its toxicity towards normal cells (11.5% viability of splenocytes at 48 h). The lowest dose of CSE, i.e. 87.6 µg/ml was also not considered since it was less effective in killing tumor cells (Fig. 2C). CSE at 146 µg/ml and 182.5 µg/ml showed 69.9% and 82.1% of EAC death at 72 h, exhibited higher killing of EAC compared to the moderate dose of BLM, i.e. 10 µg/ml being 57.2% of EAC cell death but not with the higher dose, i.e. 20 µg/ml which showed 84.7% of EAC death at 72 h (Fig. 2D). Among the two CSE doses, 182.5 µg/ml (82.1% at 72 h) showed significantly better response than 146 µg/ml (69.9% at 72 h).

### CSE treatment scavenges DPPH generation, inhibits lipid peroxidation and protects ascorbyl radical-induced protein damage

The free radical scavenging property of CSE was compared with a well known antioxidant, i.e. gallic acid (GA) in an in vitro free radical scavenging assay which generates DPPH (2,2-diphenyl-1-picrylhydrazyl). The percentage of DPPH scavenging increases gradually with increase in the concentration of CSE (49.8% at 87.6 µg/ml, 70.5% at 146 µg/ml, 80.2% at 182.5 µg/ml and 83.9% at 219 µg/ml). All doses of CSE showed significantly higher scavenging percentage over the control (EtOH). The percentage of scavenging with 219 µg/ml of CSE showed the maximum scavenging percentage, i.e., 83.9% which was comparable to GA (90.5%) (Fig. 3A).

**Figure 3.**
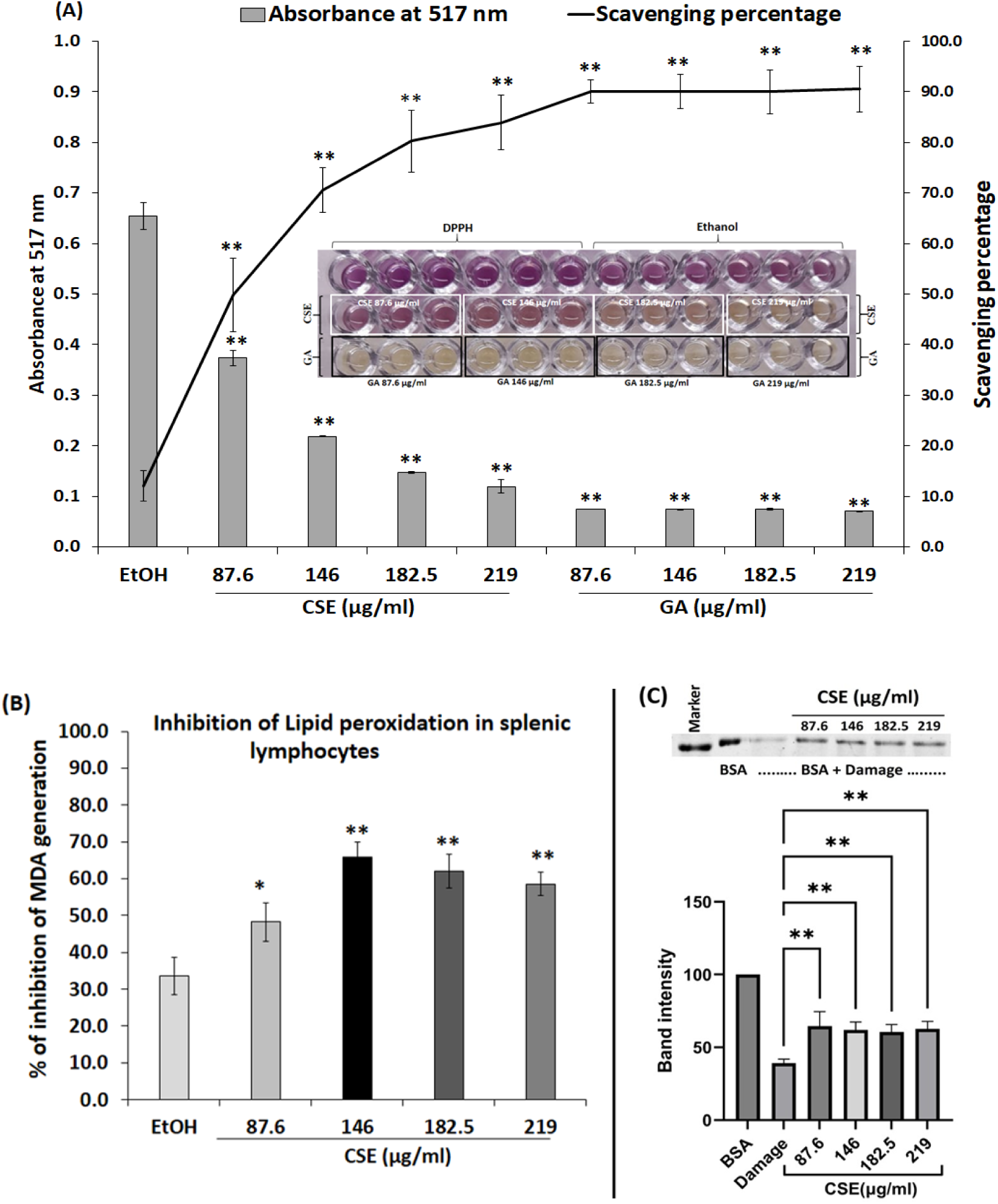
Free radical scavenging property of CSE. **A)** DPPH scavenging activity of CSE and Gallic acid (GA) at various dose concentrations (87.6 µg/ml, 146 µg/ml, 182.5 µg/ml and 219 µg/ml). The X axis depicts the concentration of CSE and gallic acid (GA) in µg/ml, Y axis depicts the absorbance of CSE, GA and ethanol (EtOH) at 517 nm and the Z axis shows the scavenging percentage of CSE and GA at various concentrations. GA has been used as a positive control. Statistical significance of CSE treatment was compared with EtOH. The photograph of the wells shows gradual change in the colour from purple to pale yellow with gradual increase in CSE dose concentration, suggesting inhibition of DPPH generation. Photograph was taken prior to the OD reading. **B)** Percentage of inhibition of lipid peroxidation of murine lymphocytes treated with different doses of CSE (87.6 µg/ml, 146 µg/ml, 182.5 µg/ml and 219 µg/ml). All doses of CSE showed free radical scavenging properties over the control. The statistical significance was calculated by comparing the EtOH treated lymphocytes with CSE-treated lymphocytes. **C)** Photograph of the coomassie stained SDS-PAGE gel shows the control protein (BSA), damaged BSA and damaged BSA with different doses of CSE (87.6 µg/ml, 146 µg/ml, 182.5 µg/ml and 219 µg/ml) taken through gel imaging system (ChemiDoc, Biorad). Band intensity was measured through Image J software and was represented in the form of bar diagram. CSE was capable of providing protection to the damaged protein with all its doses. Statistical significance was compared between the band intensity of damaged BSA with CSE-treated damaged BSA. For each graph, all significant values were marked with an asterisk (\**P* < 0.05, \*\**P* < 0.01, \*\*\**P* < 0.001 and ns; insignificant). Data are represented by the mean ± standard deviation of triplicate values for the DPPH scavenging assay and lipid peroxidation assay. For protein damage assay data are represented as mean ± standard deviation of three different gels.

Next, we wanted to examine whether CSE had a cytoprotective property by inhibiting free radical generation. CSE inhibited ascorbate-induced lipid peroxidation in murine splenic lymphocytes by quenching free radical generation. Lipid peroxidation in splenic lymphocytes results in malonaldehyde (MDA) production. The MDA production in splenic lymphocytes was inhibited by different doses of CSE (48.2% at 87.6 µg/ml, 65.9% at 146 µg/ml, 62.1% at 182.5 µg/ml and 58.5% at 219 µg/ml) (Fig. 3B).

The capability of CSE to protect protein from free radical generation was also examined. The results showed that the damage to bovine serum albumin (BSA) induced by copper ascorbate system was recovered in presence of different doses of CSE (Fig. 3C). The intensity of coomassie blue stained BSA protein band in absence of the damage system was considered as 100%. The intensity of the band falls to 39.2% after incubation with the damage system, confirming damage to the protein (BSA). With 87.6 µg/ml, 146 µg/ml, 182.5 µg/ml and 219 µg/ml of CSE treatment, the band intensity was 64.6%, 61.9%, 60.5% and 62.7% respectively (Fig. 3C). The damaged BSA protein containing different doses of CSE showed significant higher band intensity than the damaged BSA, suggesting rendering protection from the damage.

### CSE causes upregulation of p53 and downregulation of Bcl-xL proteins in EAC

CSE was found to be inhibitory towards tumor cells (EAC) proliferation and non-toxic towards normal cells (murine lymphocytes). As mentioned earlier, the lowest (87.6 µg/ml) and the highest dose (219 µg/ml) were not considered for further experiments. Two doses of CSE (146 µg/ml and 182.5 µg/ml), which showed comparable results with UNT group (control) for splenic lymphocyte growth and also exhibited significant killing of tumor cells were chosen. To confirm whether CSE induced apoptosis in tumor cells, we decided to examine two key apoptosis-related proteins p53, a pro-apoptotic protein, and Bcl-xL, an anti-apoptotic protein. The protein expression of these two proteins were examined via western blot (WB). Following 24 h treatment with CSE (146 µg/ml), EAC showed significant upregulation of p53 protein (Fig. 4A and 4B) compared to UNT (control), whereas the expression of anti-apoptotic protein Bcl-xL (Fig. 4C and 4D) was significantly downregulated in EAC compared to the control (UNT), similar to the effect brought about by 20 µg/ml of BLM; however, the results with BLM were not significant in case of Bcl-xL (Fig. 4C and 4D).

**Figure 4.**
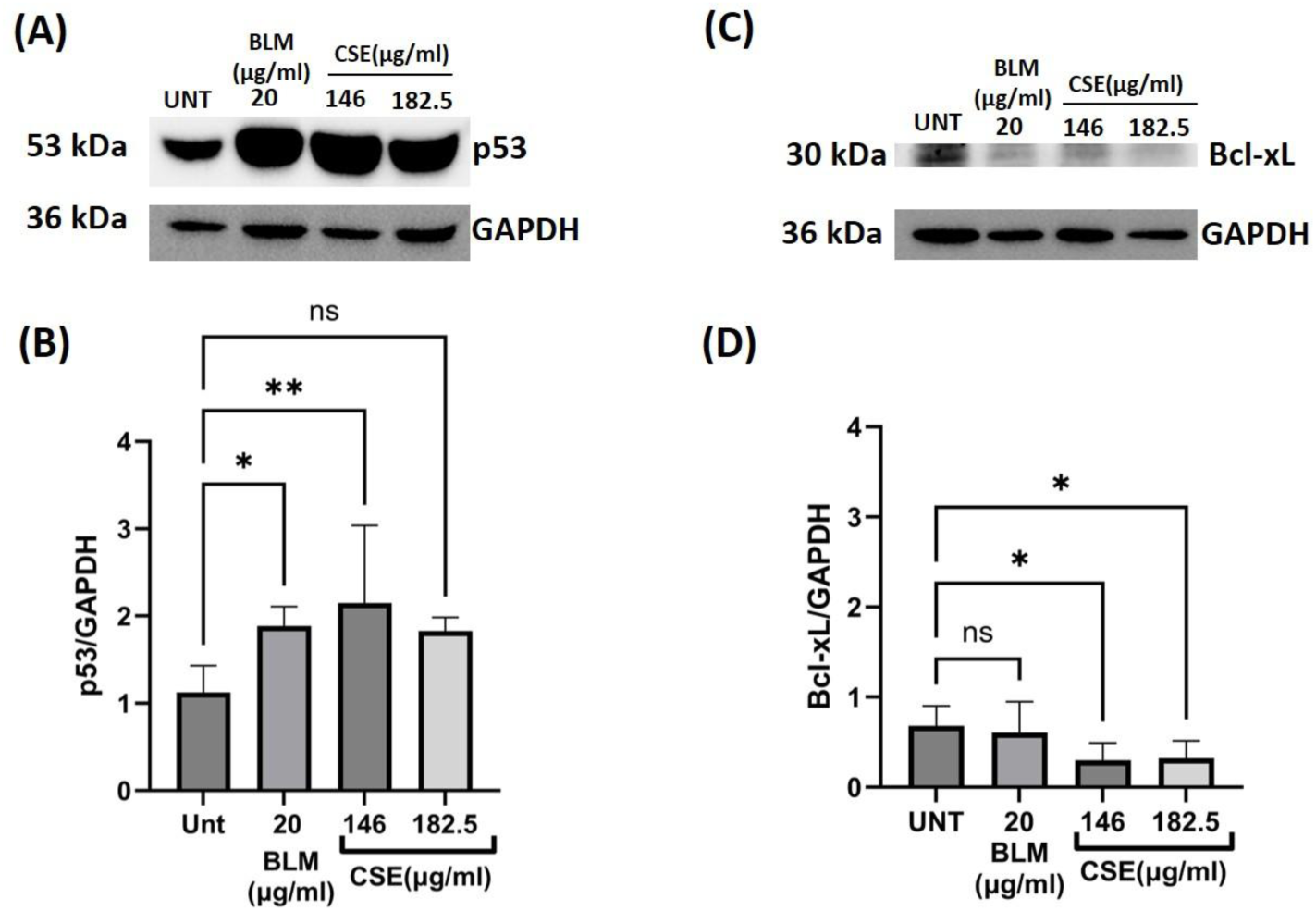
Western Blot analysis of protein isolated from EAC cells were treated with CSE in vitro (146 µg/ml and 182.5 µg/ml) and bleomycin (20 µg/ml) for 24 h. **A)** Image of the blot for p53 and GAPDH, **B)** graphical representation of the band intensity of p53 *vs* GADPH, **C)** Image of the blot for Bcl-xL and GAPDH & **D)** graphical representation of the band intensity of Bcl-xL *vs* GADPH. Bar diagram represents the mean ± SD of three independent blots. Relative protein level was estimated compared to GAPDH. Western blot analysis revealed significant upregulation of pro-apoptotic proteins p53 and downregulation of anti-apoptotic protein Bcl-xL with CSE treatment. The statistical significance of protein expression in the CSE treated groups was compared with the untreated control and was marked with an asterisk (\**P* < 0.05, \*\**P* < 0.01, \*\*\**P* < 0.001 and ns; insignificant). Both photographs were taken through gel imaging system (ChemiDoc, Biorad). Image J software has been used to determine the band intensity.

### CSE induces apoptosis-associated changes in EAC cells

Apoptotic programming leads to certain specific morphological changes in the cell. Thus, EAC cells were treated with CSE for 24 h and then stained with Giemsa; we observed under bright field microscope morphological changes consistent with apoptosis, showing surface blebbings (temporary protrusions) and formation of apoptotic bodies (Fig. 5B). However, the untreated control cells (UNT) retained the plasma membrane integrity with no blebbings (Fig. 5A). Scanning electron (SEM) and transmission electron microscopy (TEM) were also performed for CSE treated (146 µg/ml) EAC cells. Under SEM, the control (UNT) EAC cells maintained a round morphology and showed evenly distributed membrane ruffles (Fig. 5C). Following CSE treatment, the EAC cell membrane showed extensive surface protrusions and depressions leading to deformation of its round shape (Fig. 5D). Under TEM, the control (UNT) EAC cells had characteristic ruffles/ microvilli-like projections off plasma membrane and an uniform distribution of chromatin in the nucleus (Fig. 5E); however, the CSE treated tumor cells lost the surface projection, and instead, surface blebbings were observed, yielding apoptotic bodies attached to the cell membrane. CSE treated tumor cells also showed chromatin condensation and likely nuclear fragmentation (Fig. 5F).

**Figure 5.**
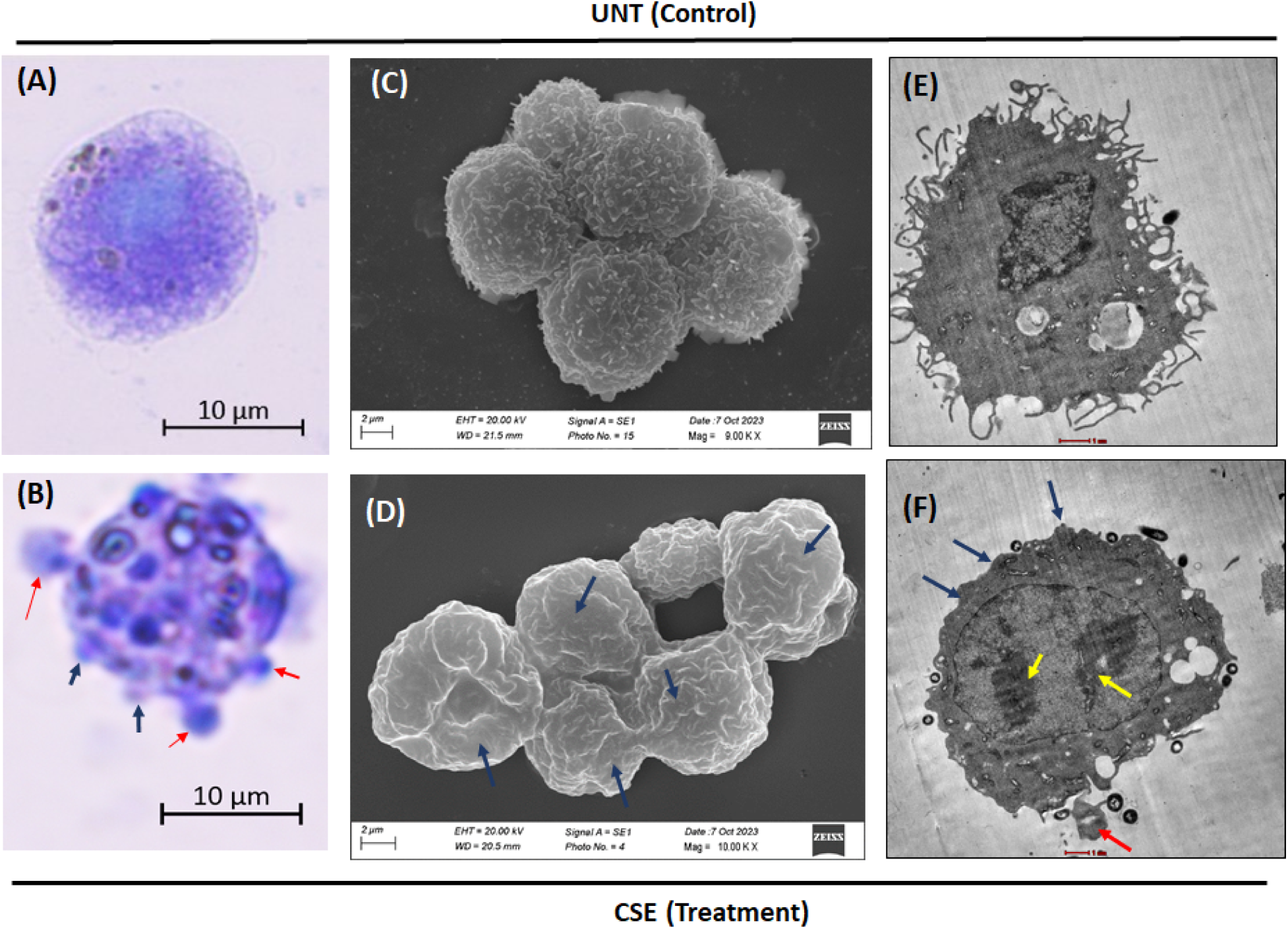
Apoptotic changes in the external and the internal milieu of EAC cells treated with CSE (in vitro) after 24 h. **A)** and **B)** giemsa stained EAC cells under brightfield microscope, **C)** and **D)** SEM images of EAC cells, **E)** and **F)** TEM images of EAC cells. Image **A), C)** and **E)** were from UNT (control) group and **B), D)** and **F)** were of CSE treated EAC cells. All EAC cells after 24 h of CSE treatment showed surface blebbings (blue arrow) and apoptotic body formation (red arrow) under brightfield microscope and under SEM. TEM images showed chromatin condensation and nuclear fragmentation (yellow arrow).

### CSE treatment induces nuclear fragmentation in EAC cells

Our in vitro experiments indicated induction of apoptotic programming and death of EAC cells following CSE treatment. Thus, to confirm the results using in vivo model, we treated the EAC tumor bearing mice intravenously with CSE, and after 24 h, collected the EAC cells from the peritoneum of the mice, and stained them with DAPI for observing nuclear changes under fluorescence microscope (Zeiss). We have published earlier that 3.65 µg/g body weight dose of CSE was non-toxic in Swiss albino mouse, hence, this dose was used for in vivo experiments ^47^. The control groups, UNT (Fig. 6A and 6B), and EtOH (Fig. 6C and 6D) showed compact nuclear morphology staining deeply with DAPI, whereas, the CSE treated cells showed several nuclear fragmentations marked with orange arrow (Fig. 6E and 6F). Nuclear fragmentation is one of the hallmarks of apoptosis; thus, it is evident that in vivo CSE treatment induced apoptotic changes in the EAC cells.

**Figure 6.**
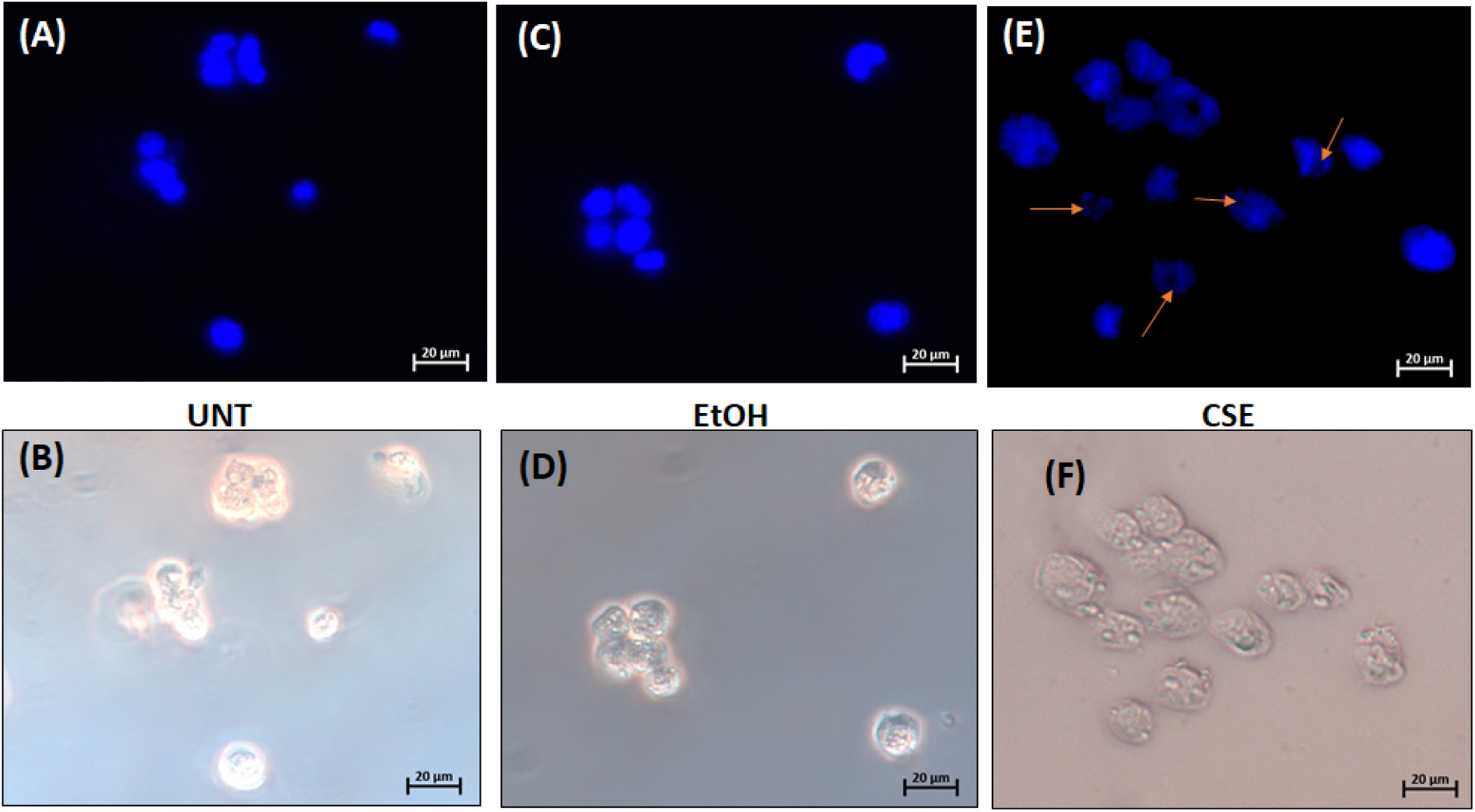
Fluorescence microscopic image of DAPI stained EAC cells collected from the peritoneum of EAC bearing Swiss albino mice. **A)** and **B)** untreated cells (control), **C)** and **D)** EtOH treated (control), **E)** and **F)** CSE (3.65 µg/g b.w.) treated Ehrlich ascitic carcinoma cell bearing mice after 24 h of treatment. **A)**, **C)** and **E)** are fluorescence microscopic images and **B)**, **D)** and **F)** are their respective brightfield microscopic images. The arrows points to the detected nuclear fragmentation with CSE treatment at 40X magnification. CSE treatment showed nuclear fragmentation of EAC cells, suggesting apoptosis

### Upregulation of p53 and downregulation of Bcl-xL proteins in vivo following CSE treatment

Fully grown ascites tumor mice (on the 20^th^ day of tumor induction) were treated intravenously (i.v.) with CSE (3.65 µg/g b.w.) and after 24 h, EAC cells from the mouse peritoneum were analyzed for the expression of pro-apoptotic protein, p53 and anti-apoptotic protein Bcl-xL through western blot. The CSE treated EAC cells showed higher level of p53 protein (Fig. 7A) and lower level of Bcl-xL protein through WB analysis (Fig. 7A) compared to UNT and EtOH treated controls. EAC cells were also stained with giemsa to observe morphological condition of the cell. Giemsa staining showed surface blebbings/protrusions (marked with blue arrow) and numerous apoptotic body formation (marked with red arrow) (Fig 7Bc) in EAC cells treated with CSE (intravenous) whereas the control cells maintained the usual round morphology (Fig. 7Ba). Under brightfield microscope the EAC cells with blebbings stained with trypan blue (Fig. 7Bd), confirming that they were not alive, whereas the control cells excluded trypan blue as their membrane was intact and thus, were not dead (Fig. 7Bb). This confirms that CSE treatment (3.65 µg/g b.w.) initiates apoptosis in EAC cells in vivo.

**Figure 7.**
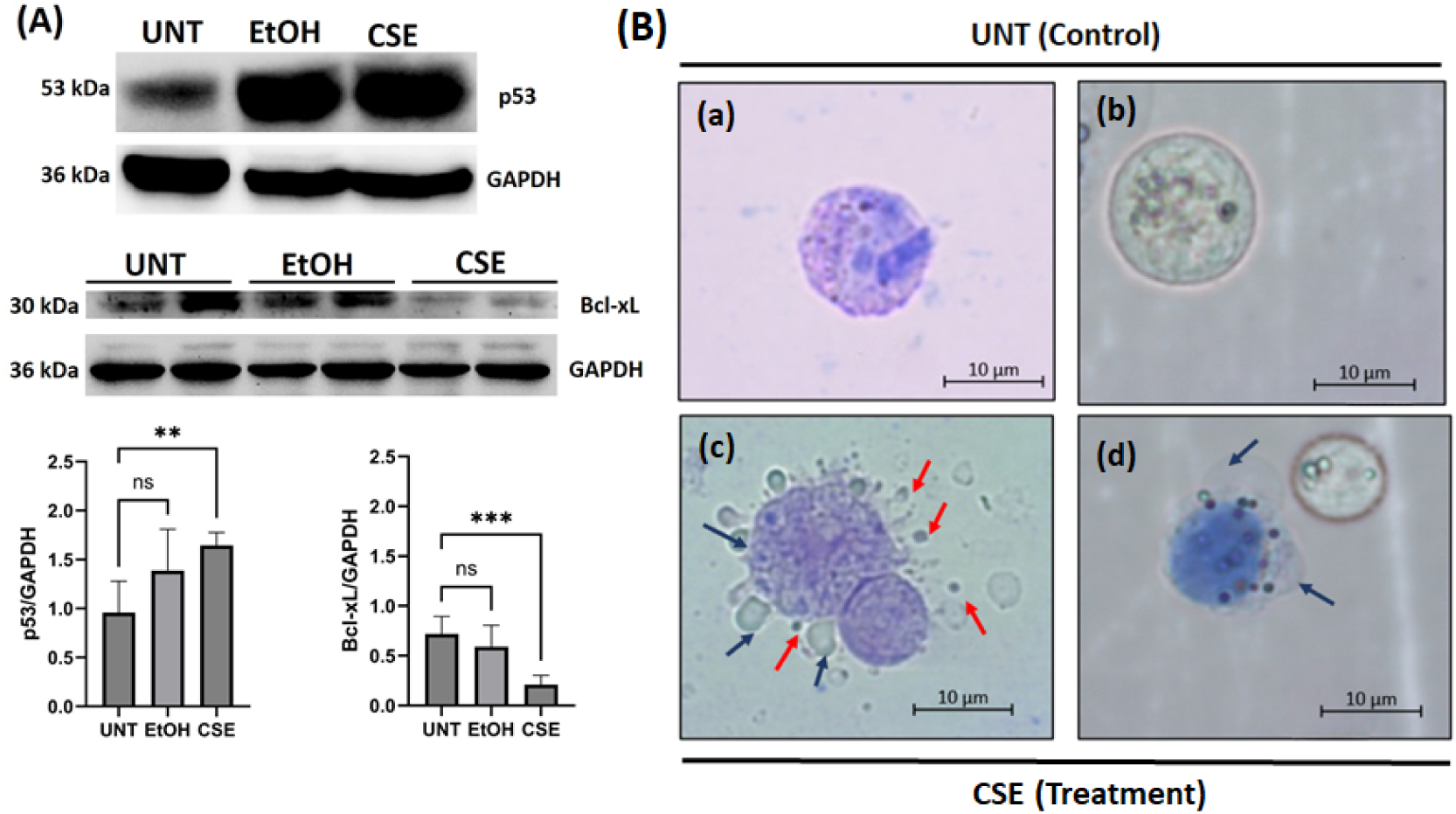
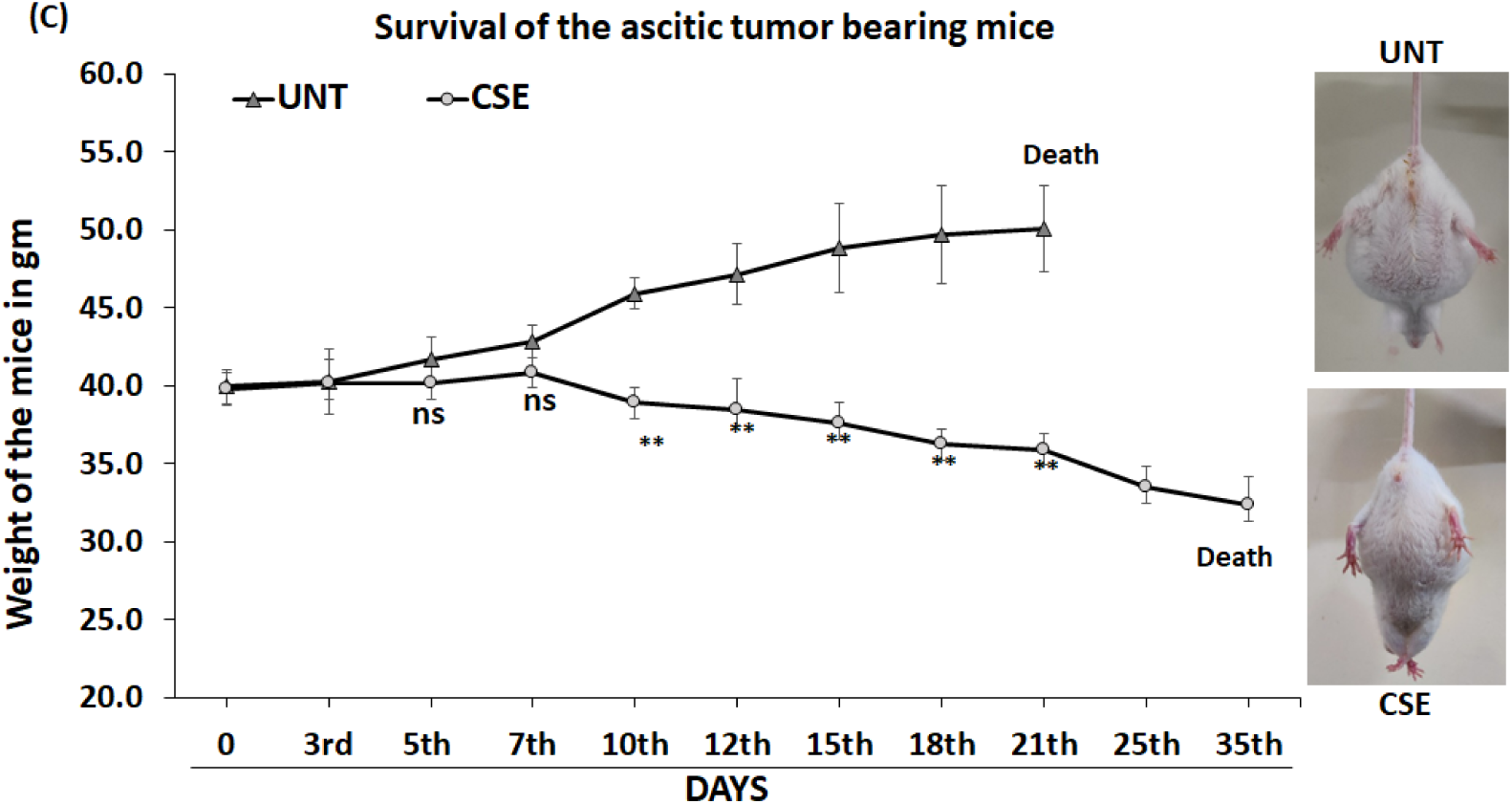
A) In vivo western blot analysis of protein expression p53 and Bcl-xL in EAC cells treated with CSE in vivo. Upregulation of p53 protein and downregulation of Bcl-xL protein was observed with CSE treatment compared to control groups (UNT and EtOH). Relative level of protein expression was estimated compared to GAPDH. Statistically significance of protein expression of the CSE treated and EtOH groups was compared with the UNT group and were marked with an asterisk (\**P* < 0.05, \*\**P* < 0.01, \*\*\**P* < 0.001and ns; insignificant). Bar diagram represents the mean ± SD of three independent blots. Image J software has been used to determine band density. **B)** EAC cells images under brightfield microscope stained with giemsa **(a)** UNT, **(c)** CSE treated and **(b)** UNT, **(d)** CSE treated were stained with trypan blue. Red Arrow indicates apoptotic bodies and blue arrow indicates surface blebbings. **C)** Rate of ascitic tumor growth with intravenous injections of CSE after 20^th^ day of ascitic tumor induction weight of mice in gm. The day of treatment is considered as Day 0 (20^th^ day of tumor induction). Statistically significant differences between the CSE treatment and UNT (control) were marked with an asterisk (\**P* < 0.05, \*\**P* < 0.01, \*\*\**P* < 0.001and ns; insignificant). CSE treatment reduces the ascites volume in EAC bearing mice as examined by measuring the body weight day wise and also restored the abdominal circumference as reflected in the photograph. **In vivo effect of CSE on ascites tumor growth and survival of mice.** As CSE was able to induce apoptosis in EAC cells both in vitro and in vivo, we expected that the treatment would likely inhibit the tumor growth and might increase the survival of the EAC tumor bearing mice. CSE (3.65 µg/g b.w.) was injected intravenously in a full grown tumor mice on the 20^th^ day of tumor induction, attaining a body weight of 40 ±1 gm and was monitored till it survives. The day of treatment was considered as day 0 (20^th^ day of tumor induction), thus, from the 21^st^ day onwards the tumor growth was monitored and the inhibition of ascites tumor growth with CSE treatment were plotted in the form of a line graph showing reduction in the ascites tumor growth and life span compared to the control. The rate of tumor development in CSE treated mice was relatively slower and survived longer than control mice (Fig. 7C). The control mice survived until 41.6 ± 3 days and the mice with CSE treatment survived for 55± 2 days (life span was calculated from the day of tumor induction till the death).

### Nature and composition of bioactive compounds present in CSE

Gas Chromatography-Mass Spectrometry (GCMS) analysis identified 43 phytocompounds, which were rich in triglyceride, fatty acid ester, alkanes, terpene alcohol, alkyl amide, fatty aldehydes, fatty amide, vitamin E, sterol, terpenoids, hydrocarbon, fatty alcohol and fatty acid (Fig. 8 and Supplementary Table 2). The major compounds were phytol (8.68%), neophytadiene (3.34%), clionasterol (3.31%), tetratetracontane (2.25%), methyl palmitoleate (1.75%), lupenone (1.55%), Palmitic acid (1.40%) cycloartenol (1.22%), sitosterone (1.12%) and 3-heptadecanol (1.06%). Several other phytocompounds such as beta-tocopherol, 2-hydroxyisocaproic acid, trimethylsilyl ester, octacosanol, triacetin, octacosane, oleamide, tetratriacontane and ricinoleic were also present.

**Figure 8.**
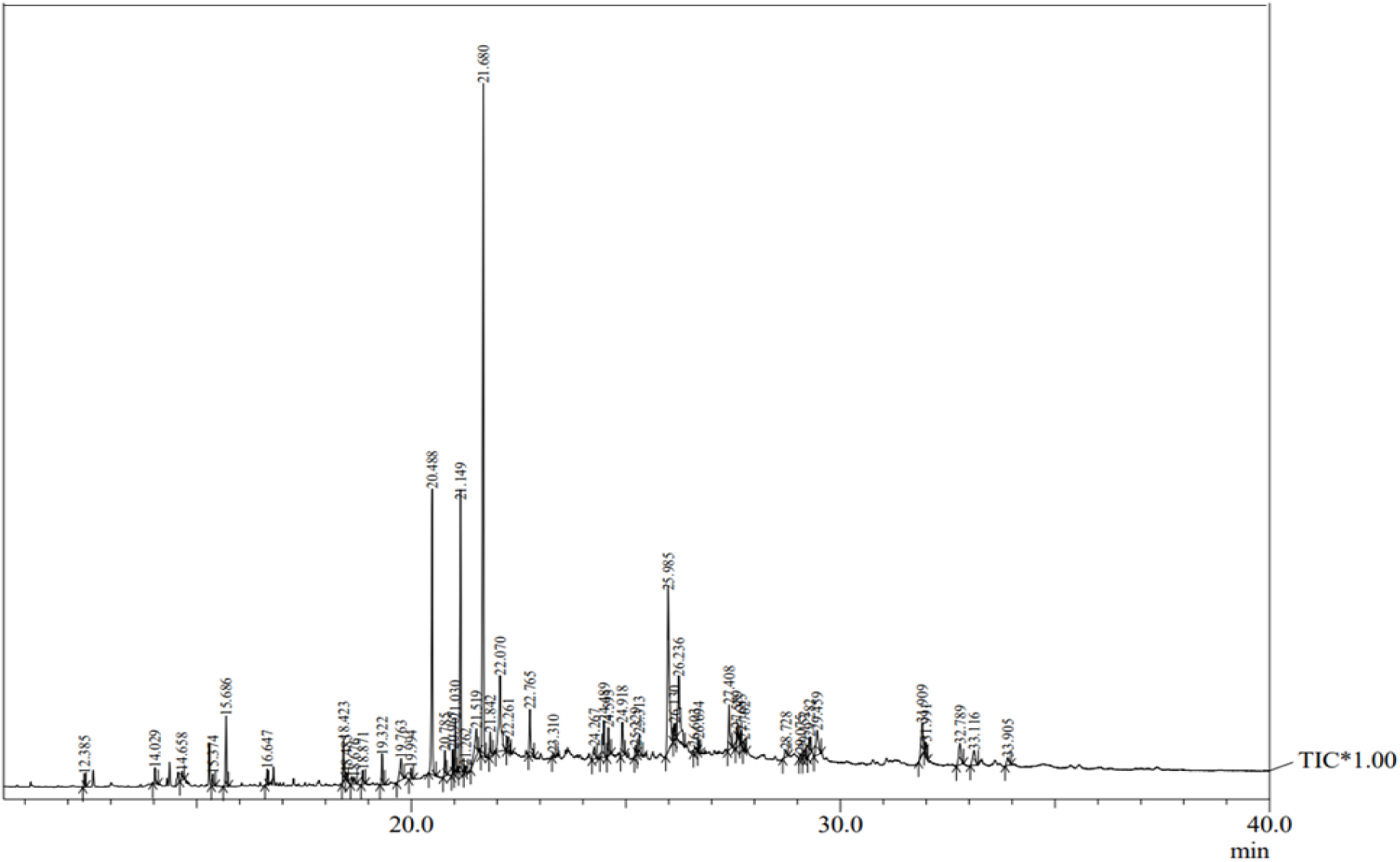
GC-MS chromatogram of the components of ethanolic leaf extract of *Cleome spinosa* (CSE). The major bioactive components are phytol (8.68%), neophytadiene (3.34%), clionasterol (3.31%), tetratetracontane (2.25%), methyl palmitoleate (1.75%), lupenone (1.55%), cycloartenol (1.22%), and 3-heptadecanol (1.06%) were the main constituents. Beta-tocopherol (0.45%), 2-hydroxyisocaproic acid, trimethylsilyl ester (0.43%), octacosanol (0.41%), triacetin (0.40%), octacosane (0.36%), oleamide (0.27%), tetratriacontane (0.23%), and ricinoleic acid (0.09%) were also traced.

### Molecular docking analysis

The present GCMS analysis and our previous study with RP-HPLC analysis ^47^ revealed several bioactive compounds in CSE. Thus, for molecular docking we selected those bioactive compounds which were present in high concentration in CSE and were also known for their anti-inflammatory and anti-tumor properties. Molecular docking was performed on the active site of the target proteins Bcl-xL and p53, focusing on the most abundant bioactive compounds identified in CSE. For molecular docking clionasterol, cycloartenol, lupenone, sitostenone, phytol, neophytadiene, oleic acid, triacetin, palmitic acid, beta-sitosterol, stigmasterol, p-coumaric acid and vanillic acid were filtered out as ligands. The compounds exhibiting the best binding affinities with both proteins were beta-sitosterol, clionasterol, cycloartenol, stigmasterol, lupenone, and sitostenone. These results are summarized in Table 1. Bcl-xL, an anti-apoptotic protein that inhibits apoptosis when overexpressed, showed promising interactions with several of these compounds. The binding affinities with beta-sitosterol, clionasterol, cycloartenol, stigmasterol, and lupenone were-9.6,-9.5,-9.4,-9.4, and-9.0 kcal/mol, respectively (Fig. 9). Beta-sitosterol demonstrated the highest binding affinity with Bcl-xL (-9.6 kcal/mol), interacting with key residues Arg139, Ala93, Ala119, Val141, and Tyr101 through alkyl and pi-alkyl bonds. Cycloartenol and stigmasterol are each bound to Bcl-xL with a binding affinity of-9.4 kcal/mol, forming conventional hydrogen bonds with Gln111 and Gln125. Clionasterol showed a binding affinity of-9.5 kcal/mol through pi-sigma, pi-alkyl, and alkyl interactions with residues Ala104, Ala119, Arg139, and Tyr101. Similarly, p53, a pro-apoptotic tumor suppressor protein, displayed strong binding affinities with these compounds. Stigmasterol bound to p53 with a binding score of-8.5 kcal/mol, forming pi-alkyl bonds with Phe212. Lupenone and sitostenone also exhibited a binding score of-8.5 and-8.3 kcal/mol, binding through alkyl bonds and van der Waals interactions involving residues Lys139, Arg267, Tyr103, Val225, and Leu264. The types of bond interaction and all the bond lengths are listed in supplementary Table 3.

**Figure 9.**
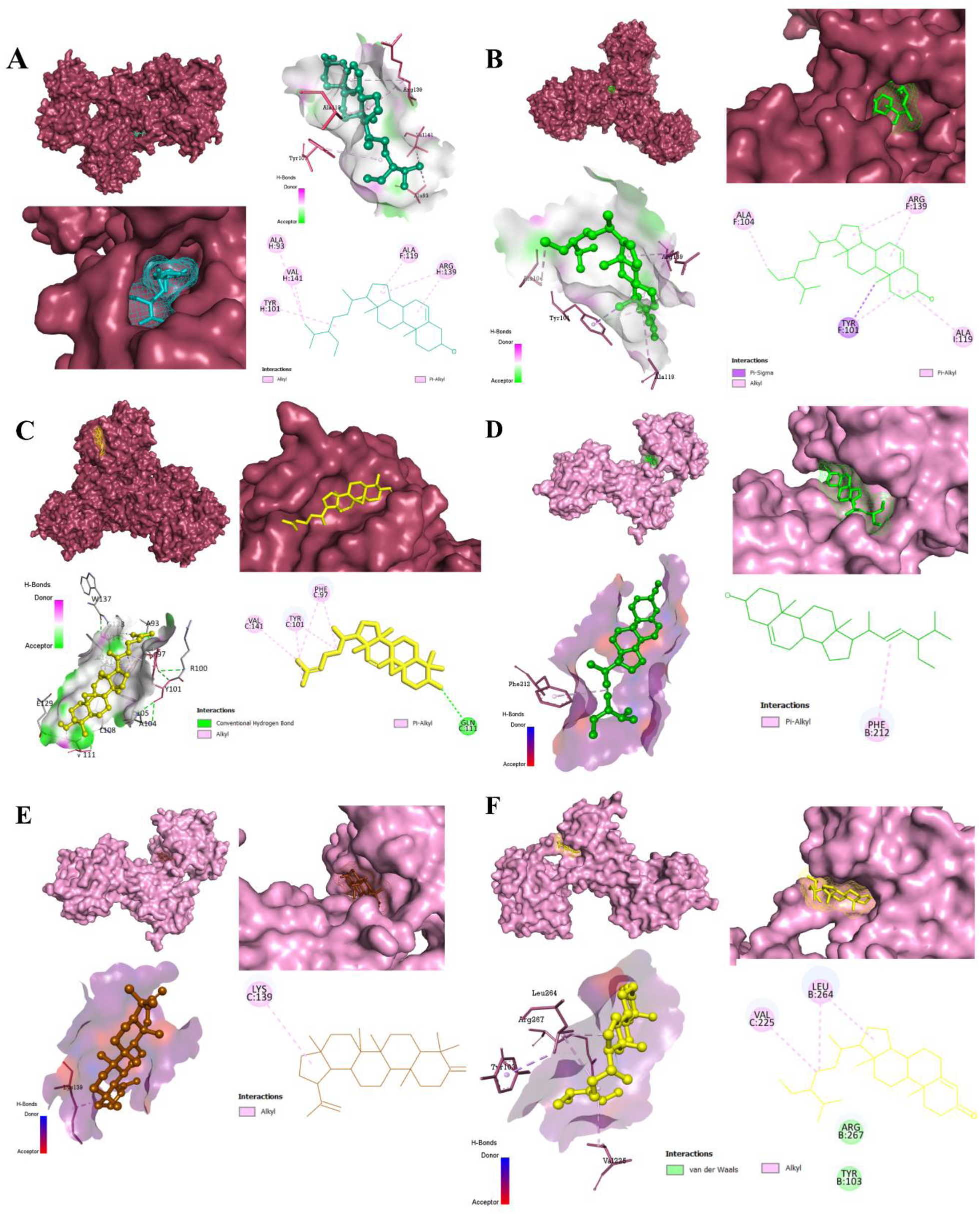
Molecular docking interaction between the CSE compounds with the target proteins Bcl-xL and p53. **A)** Binding between Beta-sitosterol and Bcl-xL with a binding score of-9.6 kcal/mol. **B)** Binding between Clionasterol and Bcl-xL with a binding score of-9.5 kcal/mol. **C)** Binding between Cycloartenol and Bcl-xL with a binding score of-9.4 kcal/mol. **D)** Binding between Stigmasterol and p53 with a binding score of-8.5 kcal/mol. **E)** Binding between Lupenone and p53 with a binding score of-8.5 kcal/mol. **F)** Binding between Sitostenone with p53 with a binding score of-8.3 kcal/mol.

**Table 1.** Binding energy scores (kcal/mol) of the Target proteins-ligands complex.

| CSE Compounds/ Ligands | Proteins |  |
| --- | --- | --- |
|  | Bcl-xL | p53 |
| Beta-sitosterol | **-9.6** | **-7.6** |
| Clionasterol | **-9.5** | **-7.9** |
| Cycloartenol | **-9.4** | **-8.7** |
| Stigma sterol | **-9.4** | **-8.5** |
| Lupenone | **-9** | **-8.5** |
| Sitostenone | **-7.9** | **-8.3** |
| Phytol | -7.1 | -4.9 |
| Neophytadiene | -6.6 | -4.9 |
| Oleic acid | -6.4 | -4.3 |
| P-coumaric acid | -6.2 | -5.9 |
| Triacetin | -5.9 | -4.8 |
| Palmitic acid | -5.9 | -4.2 |
| Vanillic acid | -5.6 | -5.5 |

## Discussion

Of the 200 species of Cleome genus, 50 are indigenous to Africa. Some of the wild species of *Cleome* are edible; its leaves, stems, pods are often consumed as vegetables due to its high vitamin and micronutrient contents in Africa ^53^. The medicinal applications of Cleome genus extend to pneumonia, diphtheria, diarrhea, malarial fever, bacterial and fungal infection, migraine, vomiting and stomach ache ^40^. The present work documented the anticancer property of *Cleome spinosa* in Ehlrich Ascites Carcinoma mouse model.

*Cleome spinosa* is an erect, foetid herb with large, beautiful deep pink flowers ^47^. It is native to new world tropics and cultivated in several parts of Asia; at some point, it must have escaped and naturalized in India. It was first identified in West Bengal in 1980 by Paria ^45^, then subsequently in Peninsular India ^54,55^. The antimicrobial, anti-inflammatory and antinociceptive activities of *Cleome spinosa* has been studied by few. Da Silva et al (2016) tested leaves and root extracts of *Cleome spinosa* with different solvents (cyclohexane, chloroform, ethyl acetate and methanol) against 12 bacterial and 5 *Candida* species. The extracts showed broad spectrum inhibitory properties against all the 17 tested bacterial and yeast species possibly due to the presence of terpenoids, flavonoid and saponins ^46^. The volatile oil extracted from the aerial part of *Cleome spinosa* were rich in oxygenated sesquiterpenes and diterpenes, likely responsible for its significant antimicrobial activity against *Streptococcus pyogenes* Group A ^56^. Methanol leaf and stem extract of *Cleome spinosa* significantly inhibited carrageenan-induced paw edema and reduced acetic acid-induced writhes in mice ^57^. We showed recently that ethanolic leaf extract of *Cleome spinosa* (CSE) significantly alleviated the deleterious DTH reaction in mouse foot pad by inhibiting TNF-α, a strong pro-inflammatory cytokine, and showed wound healing property ^47^. We also showed CSE stimulated antibody production in Swiss albino mice when challenged with sheep RBC ^48^. RP-HPLC (reversed phase-HPLC) identified several phenols, flavonoids, unsaturated hydrocarbons as well as steroids in CSE, possibly contributing to its strong anti-inflammatory and immunomodulatory properties ^47^. In this study, the anti-tumor effect of CSE was examined using EAC cell line, for the first time. Thus, we choose this plant that has never been tested for its anticancer property and is enormously rich in bioactive compounds. The animal model chosen was EAC, an undifferentiated adenocarcinoma lacking H2 antigens, and hence, shows rapid proliferation and is highly metastatic ^49,50,58,59^. It is an easily transplantable tumor model adapted to ascites form, homogeneous free tumor cells, where both inoculated number and the growth of the tumor cells can be monitored ^49,60^. These characteristics make EAC suitable for chemotherapeutic studies.

CSE significantly induced higher percentage of EAC death in vitro compared to the control groups (UNT, EtOH) in the current study. CSE exhibited dose-dependent effect; a gradual increase in the dose had a corresponding effect on EAC cell death. To know the mechanism of cell death, we examined the level of p53 protein, a potent tumor suppressor that can cause cell cycle arrest and apoptosis of tumor cells. In tumor cells, either p53 gene is mutated or its pathway is inhibited ^61^.

Using EAC, a murine mammary adenocarcinoma with the wild type p53 cell ^51,52^, we examined whether CSE can exert its anti-tumor property by activating p53. The 146 µg/ml in vitro dose and 3.65 µg/g b.w. in vivo dose (i.v.) of CSE significantly upregulated the expression of p53 apoptotic protein in our study. Interestingly, in the same tumor cells, these doses of CSE, consistently downregulated the expression of Bcl-xL, an anti-apoptotic protein. Microscopic observation confirmed the morphological alterations, typical of apoptosis such as surface blebbing, chromatin condensation, nuclear fragmentation and apoptotic body formation in the EAC cells induced by CSE in vitro as well as in vivo. CSE treatment also reduced the growth of ascites tumors and improved the survival of the tumor bearing mice.

ROS generation is one of the crucial inducers of tumorigenesis; it hyper-activates the cell signaling pathway promoting tumor cell proliferation and survival. ROS generation is heightened in tumor cells that favors angiogenesis and epithelial-to-mesenchymal transition (EMT), a major event for metastasis ^62^. At the same time, oxidative stress in healthy cells can promote carcinogenesis by causing mutations in DNA, resulting in genomic instability. Thus, maintaining ROS balance can avoid metabolic anomalies and can inhibit carcinogenesis ^63^. Antioxidant compounds are a good source of reducing oxidative stress. In the present study, CSE appeared to quench free radicals by inhibiting DPPH generation, inhibited lipid peroxidation in splenic lymphocytes, as well as recovered protein damage induced by ascorbyl radicals. We have already reported that CSE is rich in phenolic compounds such as p-coumaric acid, trans cinnamic acid, vanillic acid, ascorbic acid and gallic acid and together with presence of natural forms of vitamin E (α, β, γ, and δ-tocopherol) and unsaturated hydrocarbons (α carotene, β carotene, γ carotene, and δ carotene); the composition possibly explains its potent antioxidant capability ^47^. GCMS analysis also identified bioactive compound that are likely responsible for antioxidant and anti-tumor activity. Different groups of chemical compounds such as triglyceride, fatty acid esters, esters, hydrocarbons, fatty alcohols, fatty acids, alkanes, terpene alcohol, alkyl amide, fatty aldehydes, fatty amide, sterols, terpenoids, and others, including vitamin E were identified in the CSE via GCMS.

Molecular docking indicated that the bioactive compounds abundant in CSE exhibited strong binding affinities to key apoptosis-regulating proteins Bcl-xL and p53. Bcl-xL is an anti-apoptotic protein; its overexpression can lead to inhibition of apoptosis, and thus, can lead to tumor progression^64^. p53 is a tumor suppressor gene that regulates the cell cycle. p53 is a pro-apoptotic protein, which initiates apoptosis and helps prevent tumors ^65^. Cycloartenol, stigmasterol, clionasterol, betasitosterol, lupenone, and sitostenone showed high binding affinities with both the proteins (Bcl-xL and p53). These compounds belong to plant sterols, plant steroids, and terpenoids, which are already reported to exhibit anticancer, antioxidant, and anti-inflammatory properties. Among them, beta-sitosterol exhibited the highest binding affinity with Bcl-xL (-9.6 kcal/mol), interacting through both alkyl and pi-alkyl bonds, indicating a stable binding conformation with key residues of the protein. Cycloartenol and stigmasterol also showed strong affinities with Bcl-xL, forming conventional hydrogen bonds that enhance binding specificity. These interactions suggest that these compounds may inhibit the anti-apoptotic functions of Bcl-xL, promoting apoptosis and potentially reducing tumor growth. Similarly, the interactions between these compounds and p53 indicate their potential to impact pro-apoptotic pathways positively. Stigmasterol, lupenone, and sitostenone all exhibited significant binding affinities with p53, with stigmasterol forming pi-alkyl bonds and lupenone and sitostenone showing interactions through alkyl and van der Waals forces. The docking results, therefore, imply that these compounds may stabilize p53 and facilitate its tumor-suppressing functions, which is particularly relevant for cancer therapies targeting apoptosis induction.

These findings align with previous studies reporting the anticancer properties of these compounds. For instance, clionasterol has been reported to downregulate mitochondrial ROS levels and upregulate apoptosis signaling pathways^66^, while cycloartenol is known for its antiproliferative activity and role in apoptosis induction by downregulating Bcl-xL and upregulating pro-apoptotic proteins like Bax^67^. Stigmasterol has shown anti-tumor effects, especially in breast cancer, by downregulating anti-apoptotic proteins (such as Bcl-xL, Bcl-2, and Mcl-xL) and inhibiting angiogenesis through TNF-α and VEGFR-2 pathways^68–70^. Its efficacy against EAC in animal models further supports its therapeutic relevance in apoptosis-focused cancer treatments^71^. Lupenone, a terpenoid with documented anticancer properties^72^, has shown therapeutic potential by downregulating anti-apoptotic genes, including Bcl-2 and caspases 3, 8, and 7, in neuroblastoma^73^. Beta-sitosterol, as a plant steroid, and sitostenone has shown similar benefits, reinforcing its potential as an anti-cancer agent by supporting apoptosis and regulating cell cycles in tumor cells^74,75^. These findings underscore that CSE-derived compounds act as apoptosis modulators. Bioactive compounds present in CSE can inhibit Bcl-xL while enhancing p53 activity presenting a potential strategy for targeting apoptosis in tumor cells.

In recent years, there has been several studies where an improved therapeutic response was evident with herbal combinations or extracts that are used by indigenous communities than a single bioactive compound. Often mixtures of bioactive phytoconstituents or plant metabolites play synergistic role in their natural combinations enhancing their effectiveness in the treatment ^16,76–80^. CSE is a complex mixture of a range of bioactive compounds that are rich in antioxidant, anti-inflammatory, immunomodulatory and anti-cancer properties. Thus, examining the synergistic effects of different bioactive compounds of CSE in various tumor microenvironments, identifying their target molecules, cellular and immunological processes that they modulate will be worthwhile to investigate.

## Conclusion

The effectiveness of CSE as an apoptosis-inducing agent for tumor cells was confirmed through significant upregulation of p53 protein and downregulation of Bcl-xL both in vitro and in vitro. CSE also reduced the ascites tumor burden and increased the survival of tumor bearing mice. Thus, a mechanistic approach into how CSE can be utilised effectively as a novel adjunct therapeutics with minimal side effects merits further research.

## Materials and Methods

### Animals

Swiss albino mice of 12-14 weeks were used for the study. Ehrlich ascites carcinoma (EAC) induced mice were obtained from Chittaranjan National Cancer Institute, Kolkata, India and maintained through serial passages. To continue with the ascites cell line, adult mice of both the sexes were injected intraperitoneal with 10^6^ EAC cells per mouse in 0.1 ml PBS. After 15-20 days, a full-blown tumor would develop. All experiments were conducted in triplicate, with 10 mice in each group. Animal experiments were approved by the Institutional Animal Ethics Committee as per CPCSEA guidelines (F.No. 25/250/2012-AWD; IAEC/CBPBU/ZOO/2022/001).

### Preparation of *Cleome spinosa* leaf extract and doses

The *Cleome spinosa* leaves were collected during the month of February and March from the same location each year. The herbarium of the plant specimen was submitted to the Department of Botany, University of North Bengal for the identification (Accession number 09874) ^48^. Collected leaves were cleaned and dried in the shade. The dried leaves were then grinded to powder and soaked in ethanol (80%v/v) in 1:10 ratio. The mixture was stirred for 3 days on a magnetic stirrer (REMI). The filtration of CSE was done first by passing it through Whatman filter paper (Grade-2) and then refiltered through 0.22µm cellulose acetate syringe (E4780-1223, Eppendorf). For preparing different concentrations of doses, the extract was evaporated to dryness under reduced pressure (Rotary vacuum, Buchi) and then dissolved in the required volume of ethanol ^47^. The dose regime of CSE considered for in vitro experiments were 87.6 µg/ml, 146 µg/ml, 182.5 µg/ml and 219 µg/ml. 5 µg/ml, 10 µg/ml and 20µg/ml doses of bleomycin (Bleocin, KD-349) were considered as a positive control^81^. For in vivo experiments, 3.65 µg/g b.w. of CSE (25µl) was administered intravenously and as the CSE was dissolved in ethanol, respective volume of ethanol (25µl) was used as vehicle control. In our previous work, 3.65 µg/g b.w. dose of CSE was found non-toxic when administered intravenously in mouse, hence the same dose was used for in vivo treatment ^47^.

### In vitro cell culture of EAC

EAC cells were extracted using syringe from peritoneal exudates of mice with ascites tumor. The collected cells were washed twice in sterile phosphate buffered saline (PBS; Gibco). Live EAC cell number was counted using a hemocytometer with trypan blue ^17^. 1 × 10^6^ cells/well in 0.2 ml of DMEM (Gibco) were transferred to a 96-well microtiter plate (Eppendorf) supplemented with 10% v/v sterile heat-inactivated goat serum, 1% v/v penicillin/streptomycin solution and 1% v/v glutamine (200 mM). Cells were grown at 37 °C with 5% CO_2_ for 24 h, 48 h and 72 h. To examine the EAC cell killing with CSE treatment, trypan blue dye exclusion ^17^ and MTT (3-[4,5-Dimethylthiazol-2-yl]-2,5 Diphenyltetrazolium Bromide, SRL) methods ^82^ were performed. Trypan blue stained dead cells were counted in a hemocytometer under 40X objective of a phase contrast microscope (Magnus). Cultured EAC cells were also stained with Giemsa for observing surface blebbing and apoptotic bodies under brightfield microscope (ZEISS Axiolab 5).

The percentage of cell death (Trypan blue dye exclusion test) was calculated

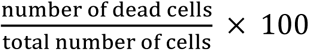

For MTT assay, tissue culture plates after different time points of treatment were centrifuged at 3000 rpm for 10 min. The supernatant was discarded and the cell pellet was washed with 1X PBS twice. MTT dissolved in PBS was added to the cell pellet at the concentration of 1 mg/ml (0.2 ml/well) and incubated in dark at 37 °C for 4 h. After incubation, the supernatant was discarded and 0.2 ml of dimethyl sulfoxide (DMSO; Rankem) was added to each well to stop the reaction. The solution was mixed thoroughly and the optical density (OD) was measured at 570 nm by a multimode plate reader (Synergy LX).

The percentage of cell death (MTT assay) was calculated as:

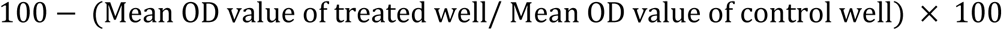

### In vitro cell culture of murine splenic lymphocytes

Spleen from normal Swiss albino mice was collected aseptically, perfused and dissociated in PBS by passing through 100 µm cell strainer (Eppendrof). After washing twice in PBS, splenocytes were centrifuged at 3000 rpm in ficoll hypaque gradient (Himedia) for 10 min. Lymphocytes were collected from the interface of ficoll hypaque and PBS, washed again twice with PBS and then adjusted to 1 × 10^6^ cells in 0.2 ml complete medium in microtiter wells. The complete medium comprised of RPMI-1640 (Gibco) supplemented with 1% v/v penicillin/streptomycin solution and 1% v/v glutamine 200 mM ^17,83,84^. Different doses of CSE (87.6 µg/ml, 146 µg/ml, 182.5 µg/ml and 219 µg/ml) were added to separate wells and incubated at 37 °C under 5% CO_2_. Cell viability was performed at 24 h and 48 h through trypan blue dye exclusion test, as previously described.

### DPPH assay

Antioxidant compounds can scavenge the DPPH (2,2-diphenyl-1-picrylhydrazyl, SRL) radical by donating hydrogen. DPPH powder was dissolved in methanol w/v to prepare DPPH solution. In glass tubes, different doses (87.6 µg/ml, 146 µg/ml, 182.5 µg/ml and 219 µg/ml) of CSE were dissolved in DPPH solution, mixed thoroughly and kept in dark for 30 min at room temperature. Finally, the solution from each tube was collected and transferred to a 96-well plate, and the absorbance of the solution was measured at 517 nm using spectrophotometry^85^. Gallic acid (Loba Chemie) was used as positive control. All tests were performed in triplicate.

The scavenging capacity was calculated as:

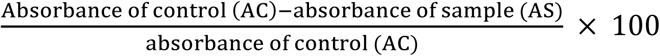

### Lipid peroxidation assay

Lipid peroxidation was induced by the copper-ascorbate system that generates malonedialdehyde (MDA) upon interaction with thioburbituric acid-reacting substances^16^. A reaction mixture was prepared by adding 20 mM Tris-HCI pH (7.4) (Himedia), 2 mM CuCl_2_ (SRL), and 10 mM ascorbic acid (SRL). The reaction mix was then added to 1×10^6^ packed splenic lymphocytes in 0.2 M sodium phosphate buffer pH (7.4) (SRL), with different doses of CSE (87.6 µg/ml, 146 µg/ml, 182.5 µg/ml and 219 µg/ml) in glass tubes and incubated for 1 h at 37° C in humidified atmosphere containing 5% CO_2_ in air. After incubation, 2 ml of TBA-TCA reagent [0.375% w/v thiobarbituric acid (TBA), 15% w/v trichloroacetic acid (TCA) and 0.25 N HCl] was added and was shaken thoroughly. The tubes were then placed in a water bath (100°C) for 15 min and then centrifuged at 1000 g for 10 min. The MDA generated in the supernatant was determined spectrophotometrically at 535 nm (U-2900, Hitachi)^86^.

The percentage of inhibition of lipid peroxidation was calculated as:

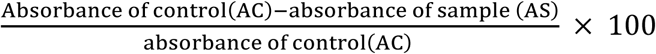

### Protein damage assay

The protein damage assay was based on a Copper-ascorbate system generating hydroxyl ion that can damage protein ^85^. The damaging system contained mixture of 100 mM sodium phosphate buffer pH (7.4) (SRL), 100-200 µM copper (II) sulfate pentahydrate (CuSO_4,_ 5H_2_O) (Loba Chemei) and 1 mM ascorbate (SRL). The damage was induced in bovine serum albumin (BSA; Himedia). In separate tubes, various amount of CSE (87.6 µg/ml, 146 µg/ml, 182.5 µg/ml and 219 µg/ml) were added and incubated for 10 min in dark. After 10 min, BSA was added to the tubes and again incubated in dark for 30 min. The control group contains only BSA dissolved in sodium phosphate buffer (pH 7.4). To stop the reaction, the tubes were placed in-80° for 15 min. Each sample were then run on reducing SDS-PAGE (10% v/v); results were analyzed by visualizing the gel in a Chemi Doc^TM^ MP Imaging System (Bio-Rad).

### Western Blot

Protein lysates were prepared from EAC cells treated in vitro (146 µg/ml and 182.5 µg/ml) and in vivo (3.65 µg/g b.w.) with CSE for 24 h. Bleomycin was used as a positive control for in vitro experiments. Proteins were estimated through Folin Lowry method and were loaded onto reducing SDS-PAGE (10% v/v). After electrophoresis, the proteins were transferred to a nitrocellulose membrane and blocked with 2% w/v BSA in TBST (20 mM Tris and 150 mM NaCl, pH 7.6 with Tween 20) or with 3% non-fatty skimmed milk in TBST (for loading control, GAPDH) for 1 h. The details of primary and secondary antibodies are listed in the supplementary Table 1. The membranes were washed followed by an overnight incubation at 4°C with primary antibody. The following day, membranes were washed with TBST and incubated with secondary antibody in TBST for 2 h at room temperature. After incubation, membranes were again washed and the protein bands were detected using immobilon western blotting substrate (Milipore, WBLUF0100) and ChemiDoc^TM^ MP Imaging System (Bio-Rad). The densitometric analysis was carried out using ImageJ software^87^.

### Scanning Electron (SEM) and Transmission Electron Microscopy (TEM)

EAC cells were treated with CSE (146 µg/ml) in vitro for 24 h, washed in PBS, and fixed for 6 h at 4 °C in Karnovsky fixative (5% glutaraldehyde and 4% paraformaldehyde in 0.1M cacodylate buffer, pH-7.4). After fixation Karnovsky solution was discarded and cells were dissolved in phosphate buffer and was sent to the Electron Microscopy Unit at the All-India Institute of Medical Sciences (AIIMS), New Delhi for further processing and imaging. For SEM, cells were imaged under scanning electron microscope, Leo 435 VP. For TEM, cells were imaged under transmission electron microscope, Philips CMIO, Netherlands^17^.

### DAPI (4’,6-diamidino-2-phenylindole) staining

EAC cells were collected from the tumor bearing mice after 24 h of CSE (3.65 µg/g b.w.) treatment intravenously, as mentioned earlier. 1 × 10^6^ EAC cells were fixed in 4% v/v formaldehyde (Himedia) for 10 min followed by three times washing in PBS and then with 0.15% v/v triton X 100 (SRL) solution for 10 min, followed by another PBS wash three times. Cells were then incubated in dark with DAPI (stock solution 1 mg/ml, SRL) staining solution for 10 min at 37 °C, followed by a subsequent wash with PBS by spinning at 1500 rpm for 5 min. The cell pellet was then re-suspended in PBS, mounted on the slides with 30% v/v glycerol (MERCK), and observed under a fluorescence microscope (ZEISS Scope.A1)^88^.

### Tumor growth monitoring

The efficacy of CSE in inhibiting the growth of tumor was also investigated. On the 20^th^ day of tumor induction, CSE was injected (3.65 µg/g b.w.) intravenously in a fully grown tumor bearing mice, attaining a body weight of 40 ±1 gm. The inhibition of tumor growth was determined by measuring the weight of mice in gm^83^.

### Gas Chromatography-Mass Spectrometry (GCMS)

The bioactive compounds present in CSE were analyzed via GCMS-QP2010 Plus (Shimadzu, Kyoto, Japan). For separation, 1.0 μl of sample was injected into Rxi-5-SIL MS capillary column (30 m X 0.25 mm ID X 0.25 µm film thickness) and run for 30 min. The phytocompounds were identified by comparing the mass spectra available with the national libraries, WILEY8.LIB and NIST14.LIB. The compound name, retention time, molecular formula, as well as structure were determined. The GCMS analysis was carried out at the Advanced Instrumentation Research Facility (AIRF), Jawaharlal Nehru University (JNU), New Delhi, India.

### Molecular Docking

Molecular docking was conducted on the most abundant compounds present in the crude CSE, which was identified via GCMS analysis and RP-HPLC ^47^. The primary focus was to understand apoptosis mechanism induced by CSE in EAC. For this present study, two target human proteins Bcl-xL and p53 were chosen.

### Target Protein preparation and Ligand preparation

The wild-type 3D structures of human Bcl-xL and p53 were obtained from the RCSB Protein Data Bank^89^ (http://www.rcsb.org) with PDB IDs 6UVF and 1TUP, respectively, in PDB format (Supplementary Fig. 1). To prepare the target proteins for docking, water molecules along with the heteroatoms were removed using Biovia Discovery studio, polar hydrogen atoms were added, and Kollman charges were applied using AutoDock tools-1.5.7 to prevent interference during docking. The compounds that were present in higher concentration in CSE were chosen as ligands for molecular docking. The 3D conformations of all the ligands were retrieved from the NCBI PubChem database in SDF format. Molecular docking was carried out using AutoDock Tools-1.5.7.

## Statistical Analysis

Data are expressed as mean ± standard deviation. Significant differences were determined by either one way ANOVA and Tukey’s multiple comparison tests or two-way ANOVA and Sidak’s multiple comparison tests using Prism software (GraphPad 10.3.0). \**P* < 0.05, \*\**P* < 0.01 and \*\*\**P* < 0.01 were considered statistically significant.

## Supplementary materials

Table 1, Table 2, Table 3 and Fig. 1.

## Acknowledgments

Authors thank all the staff of the Electron Microscopic Unit of AIIMS, New Delhi and thank Dr. Ajai Kumar, from AIRF of JNU, New Delhi for analyzing the GCMS results.

## Author Contribution PR

Methodology, investigation, writing-original draft, writing – review & editing, and formal analysis. **SR**: Methodology, investigation, writing – original draft, writing – review & editing, and formal analysis. **SB, TM, AS** and **SCD**: Methodology, writing – review & editing, and validation. **HY** and **UK**: Conceptualization, investigation, writing – original draft, writing – review & editing, methodology, project administration, resources, funding acquisition, and supervision. All authors read and approved the submitted version.

