## Supplementary tables and figure for "Ethanolic leaf extract of *Cleome spinosa* induces apoptosis in Ehrlich ascites carcinoma cells through upregulation of p53 and down-regulation of Bcl-xL in vitro and in vivo"

### SUPPLEMENTARY TABLES & FIGURES

#### Supplementary Table 1

| Protein | Resolving Gel % | Blocking agent with % and buffer | Primary Antibody | Secondary Antibody |
|---|---|---|---|---|
| p53 | 10% | 2% BSA in TBST | mouse monoclonal antibody (Santa Cruz Biotechnology, **1801**: sc-98, 1:1000 dilution in 1% BSA in TBST) | HRP-conjugated goat anti-mouse IgG (H+L) (ABclonal AS003) 1:2000 dilution in TBST |
| Bcl-xL | 10% | 2% BSA in TBST | rabbit polyclonal antibody (Santa Cruz Biotechnology, sc-7195, 1:1000 dilution in 1% BSA in TBST) | HRP-conjugated goat anti-rabbit IgG (H+L) (ABclonal AS014) 1:2000 dilution in TBST |
| GAPDH | 10% | 3% non-fatty skimmed milk in TBST | mouse monoclonal antibody (ABclonal AC033,1:50000 dilution in TBST) | HRP-conjugated goat anti-mouse IgG (H+L) (ABclonal AS003) 1:5000 dilution in TBST |

#### Supplementary Table 2

| Molecular name | Retention Time (RT) | Area | Area% | Mol. Weight (Da) | Mol. formula |
|---|---|---|---|---|---|
| Triacetin | 12.385 | 193430 | 0.40 | 218 | $C_9H_{14}O_6$ |
| Allyl heptanoate | 14.029 | 257179 | 0.53 | 170 | $C_{10}H_{18}O_2$ |
| Ethyl palmitate | 19.994 | 166855 | 0.35 | 284 | $C_{19}H_{38}O_2$ |
| Methyl Undecenate | 22.765 | 969315 | 2.01 | 198 | $C_{12}H_{22}O_2$ |
| Methyl linolelaidate | 20.967 | 343509 | 0.71 | 294 | $C_{19}H_{34}O_2$ |
| Methyl Palmitoleate | 24.489 | 843003 | 1.75 | 268 | $C_{17}H_{32}O_2$ |
| 2-Palmitoylglycerol | 24.593 | 528396 | 1.10 | 330 | $C_{19}H_{38}O_4$ |
| cis-13-Docosenoic acid, trimethylsilyl ester | 25.313 | 474725 | 0.98 | 410 | $C_{25}H_{50}O_2Si$ |

| | | | | | |
|---|---|---|---|---|---|
| Methyl triacontanoate | 29.459 | 946967 | 1.96 | 466 | $C_{31}H_{62}O_2$ |
| 2-Hydroxyisocaproic acid, trimethylsilyl ester | 14.658 | 208552 | 0.43 | 204 | $C_9H_{20}O_3Si$ |
| Diethyl phthalate | 15.686 | 1041044 | 2.16 | 222 | $C_{12}H_{14}O_4$ |
| Methyl oleate | 21.030 | 755669 | 1.57 | 300 | $C_{19}H_{36}O_2$ |
| Neophytadiene | 18.423 | 636754 | 3.34 | 278 | $C_{20}H_{38}$ |
| Tetratetracontane | 25.985 | 4763142 | 2.25 | 618 | $C_{44}H_{90}$ |
| Distearyl Phosphate | 21.680 | 10750891 | 22.29 | 602 | $C_{36}H_{75}O_4P$ |
| Octacosane | 26.694 | 171954 | 0.36 | 394 | $C_{28}H_{58}$ |
| 1-eicosanol | 18.485 | 156584 | 0.32 | 298 | $C_{20}H_{42}O$ |
| 3-Heptadecanol | 26.130 | 293645 | 1.06 | 256 | $C_{17}H_{36}O$ |
| Octacosanol | 24.593 | 528396 | 0.41 | 410 | $C_{28}H_{58}O$ |
| Methyl palmitate | 19.322 | 438874 | 0.91 | 270 | $C_{17}H_{34}O_2$ |
| Palmitic acid | 19.763 | 677171 | 1.40 | 256 | $C_{16}H_{32}O_2$ |
| Hexadecanoic acid, trimethylsilyl ester | 20.488 | 4488432 | 10.8 | 328 | $C_{19}H_{40}O_2Si$ |
| Methyl Arachate | 21.262 | 109804 | 0.23 | 326 | $C_{21}H_{42}O_2$ |
| Oleic acid | 21.519 | 1118812 | 2.32 | 282 | $C_{18}H_{34}O_2$ |
| Oleic Acid, Trimethylsilyl Ester | 22.070 | 2579580 | 5.35 | 354 | $C_{21}H_{42}O_2Si$ |
| Trimethylsilyl stearate | 22.261 | 389668 | 0.81 | 356 | $C_{21}H_{44}O_2Si$ |
| Ricinoleic acid | 23.310 | 45123 | 0.09 | 298 | $C_{18}H_{34}O_3$ |
| 10-Undecylenoyl chloride | 20.785 | 410850 | 0.85 | 202 | $C_{11}H_{19}ClO$ |
| Nonadecane | 25.229 | 168055 | 9.88 | 268 | $C_{19}H_{40}$ |
| Tetratriacontane | 29.075 | 109820 | 0.23 | 478 | $C_{34}H_{70}$ |
| Phytol | 18.423 | 636754 | 8.68 | 296 | $C_{20}H_{40}O$ |
| Decylamide | 21.842 | 367397 | 0.76 | 171 | $C_{10}H_{21}NO$ |

| | | | | | |
|---|---|---|---|---|---|
| cis-9-Octadecenal | 24.267 | 361791 | 0.75 | 266 | $C_{18}H_{34}O$ |
| Oleamide | 26.603 | 131817 | 0.27 | 281 | $C_{18}H_{35}NO$ |
| beta-Tocopherol | 28.728 | 215008 | 0.45 | 416 | $C_{28}H_{48}O_2$ |
| Clionasterol | 29.282 | 396166 | 3.31 | 414 | $C_{29}H_{50}O$ |
| Cycloartenol | 33.116 | 589784 | 1.22 | 426 | $C_{30}H_{50}O$ |
| Lupenone | 32.789 | 746657 | 1.55 | 424 | $C_{30}H_{48}O$ |
| 3,4-Dimethylphenoxyacetic acid | 15.374 | 207357 | 0.43 | 180 | $C_{10}H_{12}O_3$ |
| Sitostenone | 33.905 | 304123 | 1.12 | 412 | $C_{29}H_{48}O$ |
| 2-Monolinoleoylglycerol trimethylsilyl ether | 26.236 | 2360938 | 4.90 | 498 | $C_{27}H_{56}O_4Si_2$ |
| (Hexacosyloxy)trimethyl silane | 27.683 | 164766 | 0.34 | 556 | $C_{29}H_{62}OSi$ |
| 2-Adamantyl acetate | 27.762 | 180772 | 0.37 | 194 | $C_{12}H_{18}O_2$ |

**Supplementary Table 3.** Molecular docking results of CSE compounds with target proteins Bcl-xL and p53, including types of binding interactions and bond lengths (Å). Lower binding scores indicate stronger ligand-protein binding affinity

| PubChem ID | Ligand | Protein | Binding score (Kcal/mol) | Type of Interaction | Bond length (Å) | Interacting residue |
|---|---|---|---|---|---|---|
| 222284 | Betasitosterol | Bcl-xL | -9.6 | Alkyl, Pi-Alkyl | 5.27, 4.00, 5.02, 5.13, 3.95, 4.30, | Alanine 93 & 119, Arginine 139, Valine 141, Tyrosine 101 |
| 457801 | Clionasterol | Bcl-xL | -9.5 | Pi-Sigma, Pi-Alkyl, Alkyl | 4.06, 5.23, 4.88, 4.94, 3.42, 4.06 | Alanine 104 & 119, Arginine 139, Tyrosine 101 |
| 92110 | Cycloartenol | Bcl-xL | -9.4 | Hydrogen bond, Pi-Alkyl, Alkyl | 5.42, 4.80, 4.66, 4.07, 3.28, 3.78 | Valine 141, Tyrosine 101, Phenylalanine 97, Glutamine 111 |
| 5280794 | Stigmasterol | Bcl-xL | -9.4 | Hydrogen bond, Pi-Alkyl, Alkyl | 3.00, 4.14, 5.13,5.04, 4.56, 4.86. | Glycine 125, Arginine 139 & 132, Alanine 119, Tyrosine 101, Phenylalanine 97 |

| | | | | | | |
|---|---|---|---|---|---|---|
| 92158 | Lupenone | Bcl-xL | -9 | Carbon Hydrogen bond | 3.54 | Arginine 100 |
| 5280794 | Stigmasterol | p53 | -8.5 | Pi-Alkyl | 5.05 | Phenylalanine 212 |
| 92158 | Lupenone | p53 | -8.5 | Alkyl | 4.83 | Lysine 139 |
| 5484202 | Sitostenone | p53 | -8.3 | Van der Waals, Alkyl | 4.40, 4.78, 4.04 | Arginine 267, Tyrosine 103, Valine 225, Leucine 264 |

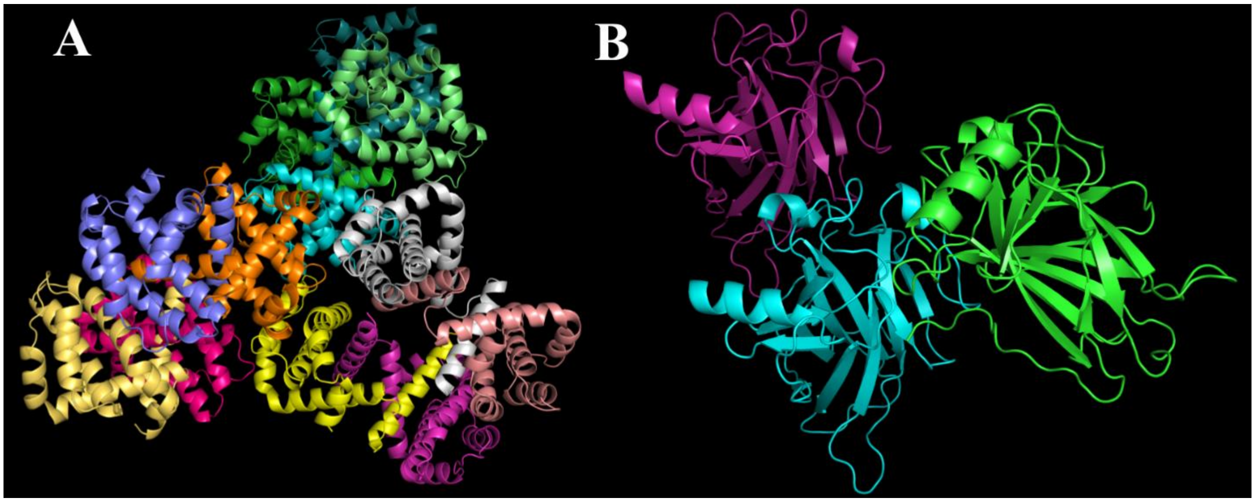


**Supplementary Figure 1.** PDB 3-dimensional structure of target proteins. (A) Bcl-xL (PDB ID: 6UVF) (B) p53 (PDB ID: 1TUP).
